# Receptor-guided engineering of oncolytic vaccinia creates a synthetic support niche for off-the-shelf NK cell therapy

**DOI:** 10.64898/2026.11.06.764990

**Authors:** Reza Rezaei, Zhiyang Gan, Ngomu Akeem Akilimali, Jordan Reff, Meiyin Lin, Yansen Xiao, Kush Parikh, Darsh Vithlani, Bakir Valentić, Alexander A Shestov, Ugur Uslu, Taha Azad, Roddy S O’Connor, Carl H. June, Neil C. Sheppard

## Abstract

Combining adoptive cell therapy with oncolytic viruses offers a means to provide localized support within solid tumors, yet viral chassis and therapeutic payloads are often selected independently of the biological state of the manufactured cell product. Here, we establish an engineering framework that uses the phenotypic and functional features of an adoptive cell product as design inputs for a complementary oncolytic niche. Using off-the-shelf expanded, frozen, and thawed (EFT) natural killer (NK) cells as a model system, we identified manufacturing-associated reductions in chemokine responsiveness, dependence on extrinsic cytokine support, and loss of sustained function at limiting effector-to-target ratios. In a head-to-head evaluation under human NK-cell antiviral pressure, oncolytic vaccinia virus (VV) retained replication and spread, whereas HSV-1 was rapidly restricted, establishing VV as a compatible viral chassis. Mapping endogenous receptor expression and applying sequential functional screening under increasingly limiting effector densities then prioritized a five-factor program comprising CXCL11, IL-15, IL-27, IL-2, and IL-1β that supported NK-cell migration, survival, cytotoxicity, and metabolic fitness at effector-to-target ratios as low as 1:10. Consolidating these signals into a single viral chassis increased antitumor activity but revealed a systemic inflammatory liability associated with constitutive IL-1β expression. We therefore placed the IL-2/IL-1β module under chemically inducible control, retaining in vivo antitumor activity while limiting systemic toxicity. Together, these findings establish a framework in which the post-manufacturing state of an adoptive cell product guides viral chassis selection, payload prioritization, and programmable control of a complementary oncolytic support niche.

**One Sentence Summary:** Cell-state-guided viral engineering sustains manufactured NK cell function at limiting effector ratios

## INTRODUCTION

Adoptive cell therapy has transformed the treatment of hematologic malignancies, largely through the success of chimeric antigen receptor (CAR) T cells (*1*), but comparable efficacy in solid tumors remains difficult to achieve (*2*). Engineering strategies have sought to address limitations of T-cell therapies (*3*), including the development of universal allogeneic products (*4*) and multitargeted constructs that reduce antigen escape (*5*). Natural killer (NK) cells possess several of these desirable features intrinsically (*6*). They can serve as readily available allogeneic effectors with a low risk of graft-versus-host disease (GvHD) (*7*), and their germline-encoded activating receptors support major histocompatibility complex-independent recognition of cellular stress and antibody-dependent cellular cytotoxicity, potentially broadening activity against heterogeneous tumors (*6*). However, the transition from fresh, feeder-expanded NK cells to cryopreserved, off-the-shelf products has revealed a substantial loss of functional potency (*8*). Unlike antigen-stimulated T cells, NK cells do not undergo robust antigen-driven clonal expansion and can become dependent on the supraphysiologic cytokine signals used during manufacturing (*9*). Current expansion and memory-induction protocols commonly use high-dose IL-2, IL-15, IL-21, IL-12, and IL-18, alone or in combination (*10–14*), and cytokine priming before cryopreservation can improve post-thaw recovery (*15*). After infusion, abrupt withdrawal of these signals can compromise persistence and sustained effector function, particularly at limiting effector-to-target (E:T) ratios, where NK cells are greatly outnumbered by tumor cells (*16*).

To improve persistence, allogeneic NK-cell products have been engineered to constitutively express autocrine survival factors such as IL-15 (*17*) or IL-21 (*18*). Although these approaches can extend NK-cell activity and have advanced into early-phase clinical evaluation (*7*), encoding a broader combination of activating cytokines directly in the cell creates a manufacturing paradox. Factors that support effector function within the tumor microenvironment may be pleiotropic or pro-inflammatory and, when expressed during ex vivo expansion, may impair cellular fitness, restrict proliferation, or induce premature differentiation (*19*). Moreover, metabolic support alone does not solve the problem of tumor access. Without directional trafficking cues, adoptively transferred NK cells may fail to localize to and penetrate solid tumors (*20, 21*). These deficiencies can be obscured in vitro by favorable E:T ratios and enforced proximity but become consequential in vivo, where relatively few NK cells must reach and control an established tumor while resisting suppressive signals and metabolites (*22, 23*). These product-specific functional requirements may therefore provide a rational set of design inputs for a localized signaling niche that coordinates recruitment, survival, and metabolic support in vivo.

Oncolytic viruses (OVs) offer a potential means of creating such a niche by selectively infecting tumors and producing immunomodulatory transgenes locally. However, because NK cells are central mediators of innate antiviral immunity (*24, 25*), combining OVs with adoptive NK-cell therapy may also create a biological conflict: activated NK cells could eliminate the vector before it establishes sufficient intratumoral spread and transgene expression. A suitable viral chassis must therefore retain tumor-selective activity and payload expression under NK-cell-mediated antiviral pressure. Among clinically developed OVs (*26*), large double-stranded DNA viruses, including adenovirus type 5 (Ad5) (*27*), herpes simplex virus type 1 (HSV-1) (*28*), and vaccinia virus (VV) (*29, 30*), provide substantial transgene capacity. Human Ad5, however, displays restricted replication in murine cells (*31*), complicating its evaluation across commonly used preclinical tumor models.

We therefore compared HSV-1 and VV to identify an oncolytic backbone compatible with adoptive NK-cell therapy. Whereas HSV-1 was rapidly restricted by activated NK cells, VV maintained viral function and spread, supporting its use as the chassis for a natural killer assistant virus (KAV). The initial KAV encoded CXCL11 to promote NK-cell migration and a sushi IL-15Rα/IL-15 complex (*32*) to support NK-cell survival and activity within the tumor microenvironment (*33*). We then used the receptor and functional state of the manufactured NK-cell product to guide cytokine selection, followed by sequential functional screening in three-dimensional tumor spheroids at increasingly limiting E:T ratios. This strategy prioritized IL-27 to enhance cytotoxic function and an IL-2/IL-1β module to support metabolic fitness under conditions of cellular stress. Because constitutive human IL-1β expression caused systemic toxicity, the IL-2/IL-1β module was placed under a vaccinia-optimized, doxycycline-inducible promoter to enable temporal control (*34*). Together, these studies establish a framework in which the biological state and functional requirements of a manufactured cell product guide the rational engineering of a complementary, programmable oncolytic support niche.

## RESULTS

### Off-the-shelf EFT NK cells exhibit product-specific and density-dependent functional deficits

Building on our observation that pre-conditioning with IL-15 and IL-18 prevents post-thaw NK cell apoptosis stemming from the leakage of granzyme B from preformed cytotoxic granules (*15*), we compared the resulting expanded-frozen-thawed (EFT) product to standard non-cryopreserved feeder-expanded (FE) (*35*) and high-dose IL-2-expanded (2E) (*36*) protocols (Fig. 1A). EFT cells demonstrated superior cytotoxicity across four solid tumor models at high effector-to-target (E:T) ratios. However, this potency advantage was significantly diminished at lower, limiting E:T ratios where killing potential mirrored that of their FE and 2E counterparts (Fig. 1B and fig. S1A). Phenotypically, EFT NK cells exhibited a highly consistent mature effector profile across donors, characterized by significant decreases in CD56 and CD8 expression accompanied by a marked increase in TIM-3 compared to FE and 2E cells (Fig. 1C and fig. S1, B and C). (*37, 38*) Transcriptomic profiling confirmed that EFT cells are distinct from both predecessors (Fig. 1, D to F), uniquely combining the heightened cell communication and cell-adhesion signaling of 2E cells with the general maturation status of FE cells. Notably, this signature was defined by a profound downregulation of chemokine receptor genes, suggesting a manufacturing-associated reduction in chemokine responsiveness (fig. S2), as previously observed for EFT NK cells (*21*).

EFT cell cytotoxicity in in vitro 2D cytotoxicity assays strongly correlates with effector cell number and remains consistent across different solid tumor lines (Fig. 1G). This dose-dependent efficacy was recapitulated in vivo following intratumoral injection into MDA-MB-231 xenografts, where increasing cell numbers dose-dependently improved tumor control with no apparent systemic toxicity (Fig. 1H and fig. S3A). These cells maintain a consistent killing potential for multiple days post-thaw, with functional activity showing a slight enhancement following the addition of low-dose IL-2 to the media (fig. S3B). While EFT NK cells remain viable in culture post-thaw, they do not undergo further expansion (fig. S3, C and D). Furthermore, at low E:T ratios, these cells are unable to sustain effector function without continuous cytokine support in 2D assays (fig. S3E). We therefore treated these product-specific deficits in trafficking potential, cytokine dependence, and low-density function as design requirements for an extrinsic viral support platform.

**Fig. 1.**
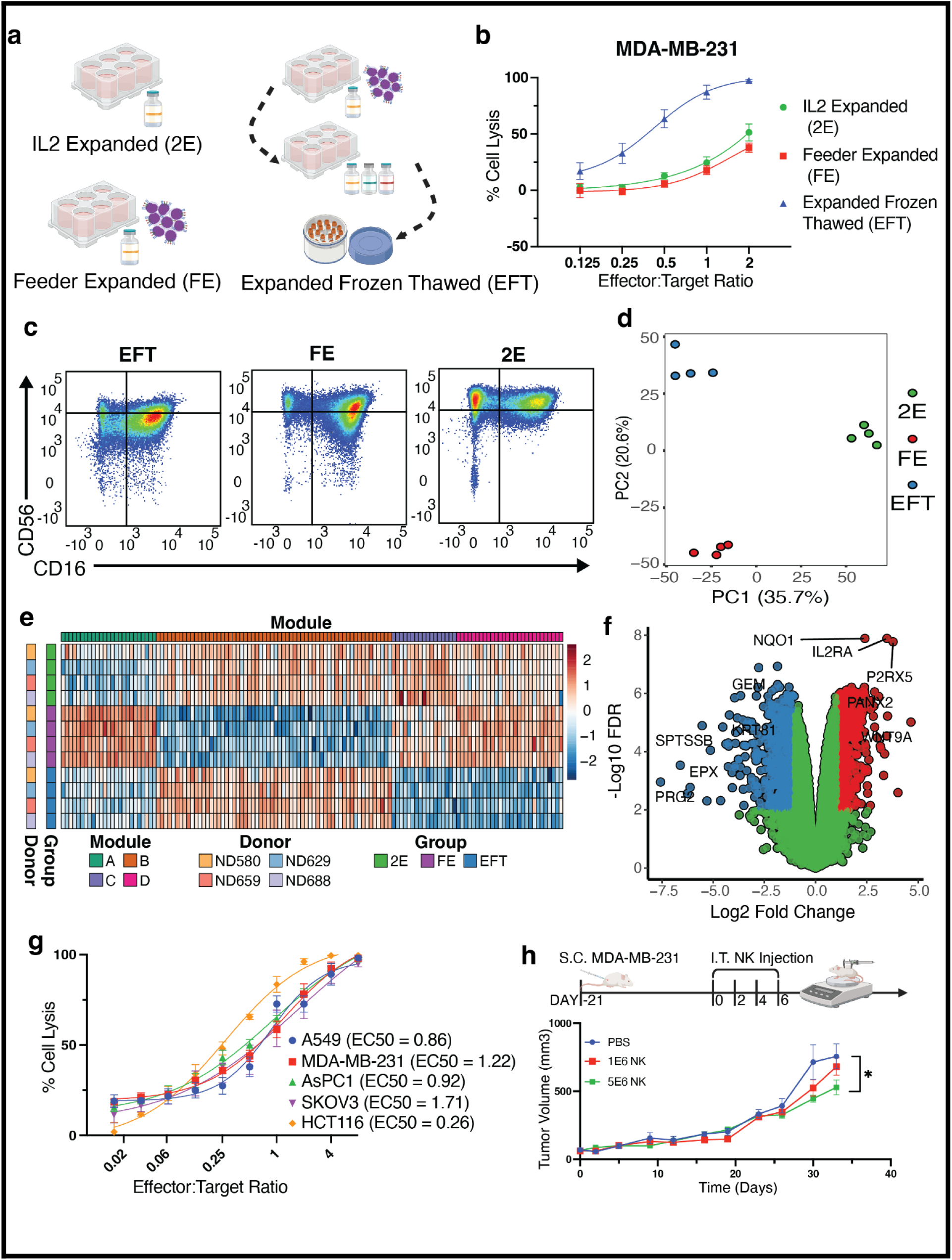
EFT NK cells are transcriptionally distinct and exhibit potent but effector-to-target ratio-dependent cytotoxicity. (A) Schematic of the manufacturing workflow for three distinct NK cell products: IL-2 Expanded (2E), Feeder Expanded (FE), and Expanded Frozen Thawed (EFT). Primary NK cells were expanded using either high-dose IL-2 (1000 U/ml; 2E) or irradiated K-IL21 feeder cells and low-dose IL-2 (100 U/ml; FE and EFT). EFT cells were activated on day 15 with a cytokine cocktail (IL-2 at 100 U/ml, IL-15 at 50 ng/ml, and IL-18 at 250 ng/ml) for 24 h prior to cryopreservation and subsequent thawing. (B) 24-hour cytotoxicity of 2E, FE, and EFT NK cells against Click Beetle Red (CBR) luciferase-expressing MDA-MB-231 cells across a range of E:T ratios (2:1 to 1:8). Percentage of tumor cell lysis was determined by luciferase signal quantification. (C) Representative flow cytometry plots showing surface expression of CD56 and CD16 on 2E, FE, and EFT NK cell products. (D) Principal component analysis (PCA) of global RNA-sequencing (RNA-seq) profiles. Percentages on the axes indicate the proportion of variance explained by PC1 and PC2. (E) Heatmap of differentially expressed genes (DEGs) in EFT NK cells compared to the mean expression of 2E and FE products. Genes are organized via hierarchical clustering into four distinct modules; gene names are omitted for illustrative clarity. (F) Volcano plot comparing the global gene expression profile of EFT NK cells against the mean of 2E and FE products. Red and blue points represent significantly upregulated and downregulated genes, respectively (adjusted P value < 0.01 and |log2 FC| > 1). The top five up-and downregulated genes are annotated. (G) Cytotoxicity of EFT NK cells against a panel of solid tumor cell lines -A549 (lung), MDA-MB-231 (breast), AsPC-1 (pancreatic), SKOV-3 (ovarian), and HCT-116 (colorectal) -across E:T ratios from 8:1 to 1:64. Curves were generated using a four-parameter non-linear regression model. The calculated EC50 (E:T ratio for 50% lysis) is reported for each line. Data are mean +/-SEM from four independent biological donors (ND607, ND608, ND410, and ND500), except for HCT-116 (n = 3; ND607, ND608, and ND410). (H) In vivo antitumor efficacy of EFT NK cells against established MDA-MB-231 tumors in NSG mice. Mice received four injections (days 0, 2, 4, and 6) of PBS, 1 x 10^6 (1M), or 5 x 10^6 (5M) EFT NK cells. The first two injections utilized donor ND659 and the final two utilized donor ND669. Tumor volume was monitored via caliper measurement. Data are mean +/-SEM (n = 5 mice per group). Unless otherwise specified, data in (B) to (F) are presented as mean +/- SEM from four independent biological donors (ND580, ND629, ND659, and ND688). P values in (H) were determined using a two-tailed unpaired t test at day 33. *P < 0.05, **P < 0.01, ***P < 0.001, ****P < 0.0001.

### Viral chassis selection identifies vaccinia as compatible with activated EFT NK cells

Because EFT NK cell efficacy is constrained by high tumor burden (resulting in low E:T ratios) and by its dependence on exogenous cytokines, we hypothesized that OVs could enhance therapeutic efficacy by debulking the tumor mass while simultaneously serving as a platform for localized cytokine delivery. However, since NK cells are primary effectors of antiviral immunity, an optimal oncolytic helper virus must be built on a backbone that retains oncolytic activity and transgene expression in the presence of an activated NK cell population. We therefore performed a head-to-head comparison of Herpes Simplex Virus-1 (HSV-1) and Vaccinia Virus (VV) to identify a chassis biologically compatible with EFT NK cell therapy. At high MOI, both viruses showed broadly comparable short-term cytotoxicity across tumor lines, although HSV-1 was less effective in MDA-MB-231 cells (Fig. 2A). At lower MOI, VV maintained efficient spread in SKOV3 and A549 cells, whereas HSV-1 showed impaired replication in MDA-MB-231 cells (fig. S4A). Combination with EFT NK cells produced only modest, cell-line-dependent improvements in 2D assays (Fig. 2B), while in 3D spheroids HSV-1 displayed faster oncolysis and stronger short-term combination activity than VV (Fig. 2C). In vivo, however, activated NK cells reduced the antitumor activity of HSV-1 in SKOV3 tumors, whereas VV maintained tumor control in the presence of NK cells (Fig. 2D). Exogenous IL-2 further improved VV plus NK-cell activity in MDA-MB-231 tumors, supporting the importance of cytokine support for the cellular component (fig. S4B).

**Fig. 2.**
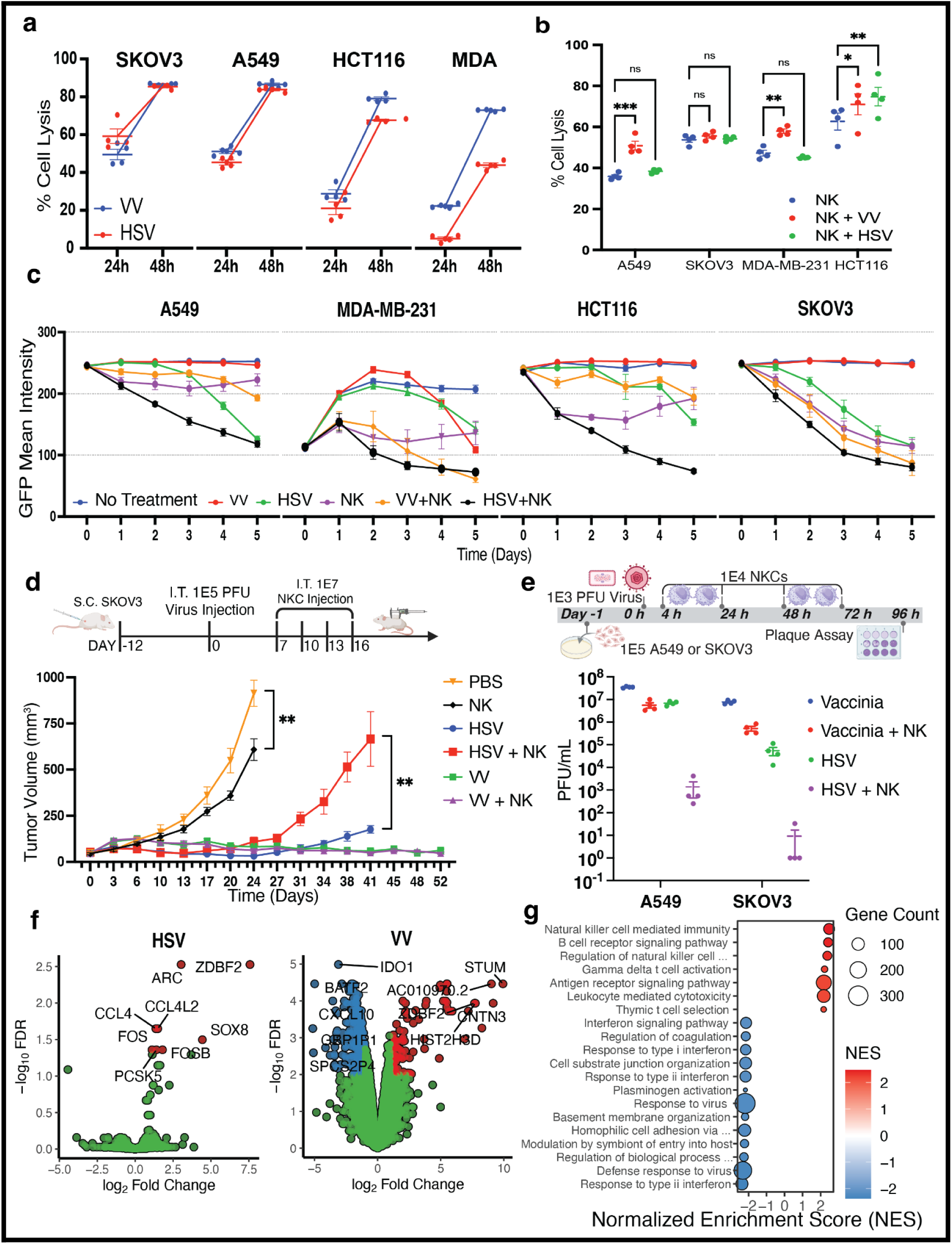
Vaccinia virus retains viral function under NK-cell pressure and supports combination therapy with EFT NK cells. (A) Comparative oncolysis of HSV (dICP6) and VV (dTK) across four solid tumor cell lines (SKOV-3, A549, HCT-116, and MDA-MB-231). Tumor cell lysis was quantified via CBR luciferase signal at 24 and 48 h post-infection (MOI 0.1). Data are mean +/- SEM from n = 4 replicates. (B) Tumor cell lysis in 2D culture across four cell lines (A549, SKOV-3, MDA-MB-231, and HCT-116) following treatment with EFT NK cells alone or in combination with VV or HSV. Cells were infected with VV or HSV (MOI 0.1) and cocultured with EFT NK cells at a 1:2 E:T ratio for 24 h. Percentage of lysis was determined by CBR luciferase quantification relative to untreated controls. (C) Longitudinal GFP mean intensity measurements in A549, MDA-MB-231, HCT116, and SKOV3 tumor spheroids following treatment with VV, HSV, EFT NK cells, or indicated combinations. Spheroids were infected (MOI 0.1) and EFT NK cells were added 4 h post-infection at a 1:2 E:T ratio. Mean intensity was monitored daily from day 0 to day 5. (D) In vivo antitumor efficacy of EFT NK cells and oncolytic viruses in a subcutaneous SKOV-3 xenograft model. NSG mice were seeded with tumor cells on day -12 and received a single intratumoral (I.T.) injection of HSV or VV (1 x 10^5 PFU) on day 0. Mice subsequently received four I.T. injections of EFT NK cells (1 x 10^7 cells per dose) on days 7, 10, 13, and 16. Tumor volume was monitored via caliper measurement. (E) Analysis of viral persistence in A549 and SKOV3 cells in the presence of EFT NK cells. Tumor cells (1 x 10^5) were infected with VV or HSV (1 x 10^3 PFU) at 0 h. EFT NK cells (1 x 10^4 per dose) were added at 4, 24, 48, and 72 h post-infection. Viral titers were determined via plaque assay at 96 h post-infection. (F) Volcano plots showing differentially expressed genes in EFT NK cells following coculture with HSV- or VV-infected SKOV3 cells. NK cells were cocultured at a 2:1 E:T ratio with tumor cells infected at MOI 0.1 and isolated 48 h post-infection for bulk RNA-seq. Plots represent the comparison of virus-infected cocultures against uninfected cocultures using the same donors. Adjusted P value < 0.05 and |log2 FC| > 1. (G) Gene set enrichment analysis (GSEA) displaying normalized enrichment scores (NES) for biological pathways significantly regulated in VV-exposed EFT NK cells compared to uninfected controls. Data in (B) are mean +/- SEM from n = 4 independent biological donors (ND602, ND607, ND669, and ND046). Data in (C) are mean +/- SEM from n = 4 independent biological donors (ND602, ND616, ND657, and ND669). Data in (D) are mean +/- SEM (n = 5 mice per group); P values determined by two-way ANOVA. Data in (E) are mean +/- SEM from n = 4 replicates. Data in (F) and (G) are from n = 3 independent biological donors (ND602, ND607, and ND657). *P < 0.05, **P < 0.01, ***P < 0.001, ****P < 0.0001.

To evaluate viral persistence in the presence of limiting NK-cell numbers, we assessed NK cell-mediated clearance at a 1:10 E:T ratio. HSV-1 was almost completely cleared by 96 hours, whereas VV titers were minimally affected (Fig. 2E). Virus-conditioned transcriptomic profiling further showed that VV, but not HSV-1, induced a marked transcriptional response in EFT NK cells, including enrichment of effector, leukocyte-recruitment, and interferon-associated programs (Fig. 2F,G and fig. S5). Together, these findings identified VV as a viral chassis capable of maintaining function under activated NK-cell pressure, supporting its selection for subsequent cell-product-guided engineering.

### Receptor-guided vaccinia engineering provides migratory and survival support to EFT NK cells

Based on its superior ability to maintain viral persistence despite NK cell-mediated antiviral pressure, we selected VV as the chassis to develop the natural killer assistant virus (KAV). To address the manufacturing-associated reduction in chemokine responsiveness identified in Figure 1, we first analyzed the expression of chemokine receptors in EFT NK cells across both resting and tumor-cocultured states (Fig. 3A). While several receptors were detected, CXCR3 exhibited the most consistent expression levels across both conditions (Fig. 3A). Flow cytometric analysis confirmed high surface expression of CXCR3, whereas CXCR4 expression was lower and donor-dependent (Fig. 3B). We therefore engineered the VV backbone to express CXCL11, a high-affinity ligand for CXCR3 (Fig. 3C). In transwell migration assays, conditioned media from VV-CXCL11-infected cells induced a significant increase in NK-cell migration compared to wild-type VV or uninfected controls, confirming the functionality of the virally encoded chemokine (Fig. 3D).

**Fig. 3.**
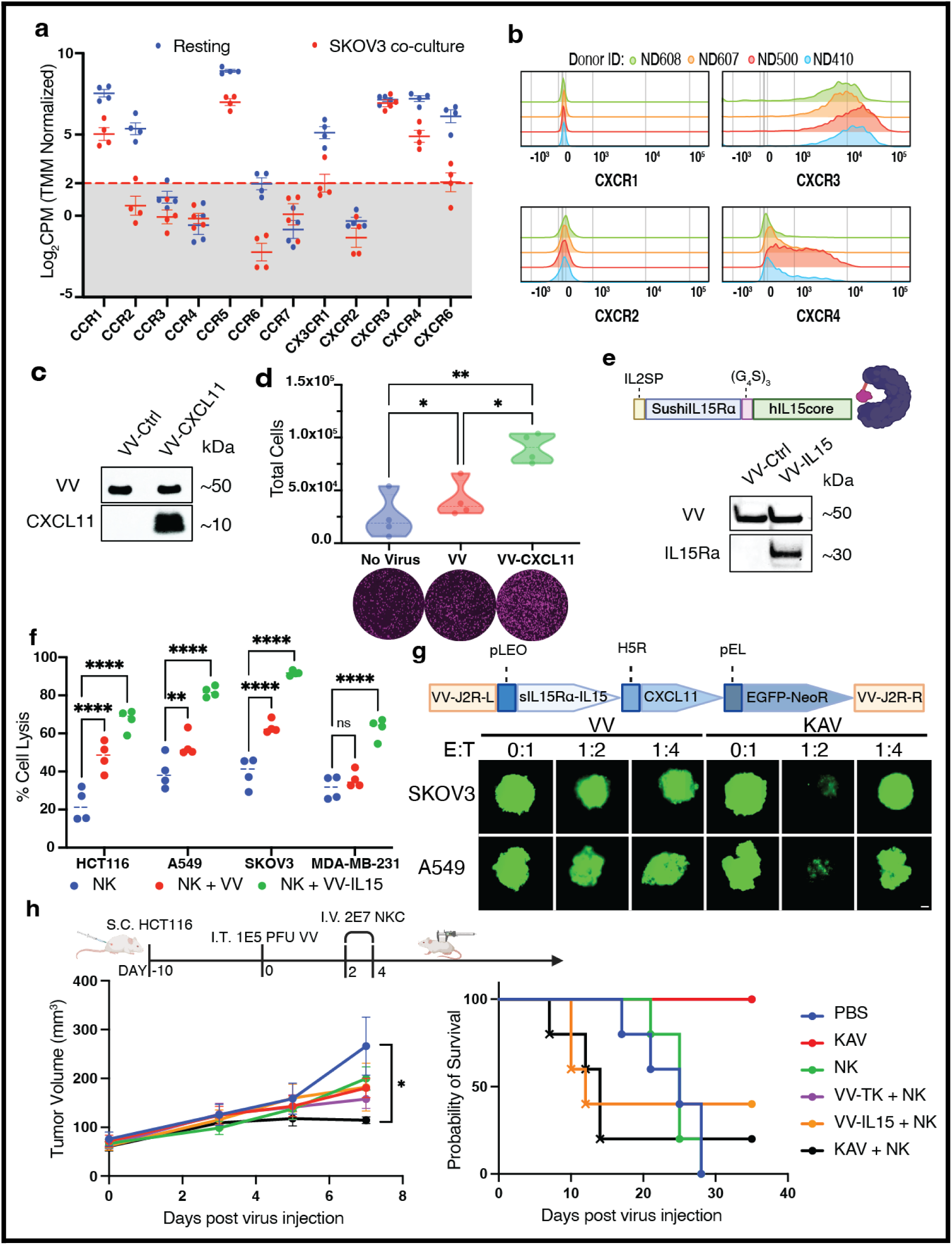
Engineering of an oncolytic vaccinia virus to enhance EFT NK cell recruitment and survival within the tumor microenvironment. (A) Chemokine receptor expression profile of EFT NK cells. Log2 counts per million (CPM) values (TMM normalized) are shown for resting (blue) and SKOV-3 co-cultured (red) EFT NK cells. Data represent n = 4 independent biological donors per condition, integrated from two independent experiments using batch effect reduction. The dashed line indicates a threshold of 2 Log2CPM. (B) Surface expression of CXCR1, CXCR2, CXCR3, and CXCR4 on EFT NK cells as determined by flow cytometry. Histograms display fluorescence intensity across four independent biological donors (ND608, ND607, ND500, and ND410). (C) Western blot analysis of CXCL11 expression in A549 cell lysates. Cells were infected with VV-Ctrl (VV-dTK) or VV-CXCL11 (MOI 0.5) and harvested 24 h post-infection. Viral D8L was used as a loading control. (D) Transwell migration of EFT NK cells toward filtered (0.2 um) supernatants from A549 cells 24 h post-infection. DiD-labeled NK cells (1 x 10^5) were incubated for 1 h in transwell inserts (5 um pore size). Top: Quantification of total migrated cells (n = 4 independent biological donors: ND616, ND502, ND657, and ND668). Bottom: Representative microscopy images of migrated cells. P values determined by one-way ANOVA. (E) Schematic and Western blot analysis of sIL15Ra-IL15 fusion protein expression in A549 cell lysates. Top: Schematic of the fusion protein construct containing the IL-2 signal peptide (IL-2SP), sushi domain of IL-15 receptor alpha (SushiIL15Ra), and hIL15 core connected by a (G4S)3 linker. Bottom: Western blot of A549 cells 24 h post-infection (MOI 0.5). Viral D8L was used as a loading control. (F) Tumor cell lysis in 2D culture across four cell lines using IL-2 deprived EFT NK cells. Cells were infected with VV or VV-IL15 (MOI 0.1) and cocultured with NK cells 4 h post-infection (1:2 E:T ratio). Percentage of lysis was determined 24 h post-treatment by CBR luciferase quantification. Data are mean +/- SEM (n = 4 independent biological donors); P values determined by two-way ANOVA. (G) Schematic and functional validation of the KAV platform in 3D tumor spheroids. Top: Schematic of the KAV construct inserted into the J2R (TK) locus, expressing sIL15Ra-IL15 (pLEO promoter), CXCL11 (H5R promoter), and EGFP-NeoR (pEL promoter). Bottom: Representative fluorescence microscopy of EGFP-expressing SKOV3 and A549 spheroids at day 5 post-treatment (MOI 0.1) at indicated E:T ratios. Data are representative of n = 4 donors. (H) In vivo antitumor activity and toxicity in an HCT116 subcutaneous xenograft model. Top: Timeline for tumor seeding (day -10), I.T. viral injection (1 x 10^5 PFU, day 0), and I.V. EFT NK cell injections (2 x 10^7 cells per dose, days 2 and 4). Left: Tumor volume measurements through day 7 post-virus injection; P values determined by one-way ANOVA on day 7. Right: Kaplan-Meier survival curves where crosses indicate toxicity-related deaths. *P < 0.05, **P < 0.01, ****P < 0.0001.

To provide localized survival support to EFT NK cells, we further engineered the virus to express a sushi-IL-15Rα/IL-15 complex (Fig. 3E), which provides enhanced signaling and stability compared to IL-15 alone (*32*). Secretion of the functional cytokine complex from infected cells was confirmed using a reporter cell line (fig. S6A). To simulate in vivo cytokine deprivation, EFT NK cells were rested for 48 hours without IL-2; under these conditions, the IL-15 complex restored cytotoxicity across all cell lines at a 1:2 E:T ratio (Fig. 3F). However, in 3D spheroid assays, complete tumor clearance was achieved at a 1:2 ratio, but efficacy was lost at a 1:4 ratio (Fig. 3G). In an intraperitoneal SKOV3 model, administration of EFT NK cells following KAV infection resulted in a transient decrease in tumor burden that eventually returned to the viral monotherapy baseline, suggesting a lack of sustained activity (fig. S6, B and C). Increasing the cell dose to overcome this threshold in a subcutaneous model failed to achieve clearance and instead induced systemic toxicity in the IL-15 combination groups (Fig. 3H). Together, these results showed that receptor-matched migratory and survival signals improved EFT NK-cell function but were insufficient at limiting effector density, while simply increasing the NK-cell dose introduced systemic toxicity.

### Sequential receptor-guided screening identifies complementary signals that sustain NK-cell function at limiting density

To overcome the functional failure observed at limiting E:T ratios, we sought to engineer a localized signaling niche tailored to the specific receptor profile of EFT NK cells. We first identified interleukin receptors with consistent expression across both resting and tumor-cocultured states (Fig. 4A). These receptor profiles were used to constrain candidate signals for engineering, and we constructed a library of nine vaccinia viruses, each encoding a distinct human interleukin, excluding IL-15 (fig. S7A). Functional activity of each virally encoded cytokine was initially verified in 2D cytotoxicity assays (Fig. 4B and fig. S7B). We next subjected this receptor-informed viral library to sequential functional screening under increasingly limiting effector densities. At a 1:5 E:T ratio in 3D tumor spheroids, KAV was combined individually with each cytokine-expressing virus (VV-Cyt), identifying IL-2 and IL-27 as the most consistent enhancers of cytotoxicity across multiple tumor models (Fig. 4C). Combinatorial screening of these prioritized signals further showed that inclusion of IL-2 and IL-27 consistently improved KAV-mediated combination activity, with additional enhancement observed when IL-1β was incorporated (fig. S8). We then increased the selective stringency to a 1:10 E:T ratio, using KAV, IL-2, and IL-27 as the new baseline for a second-stage screen. Under these conditions, IL-1β provided the most consistent additional support for NK cell-mediated tumor lysis (Fig. 4D and fig. S9).

**Fig. 4.**
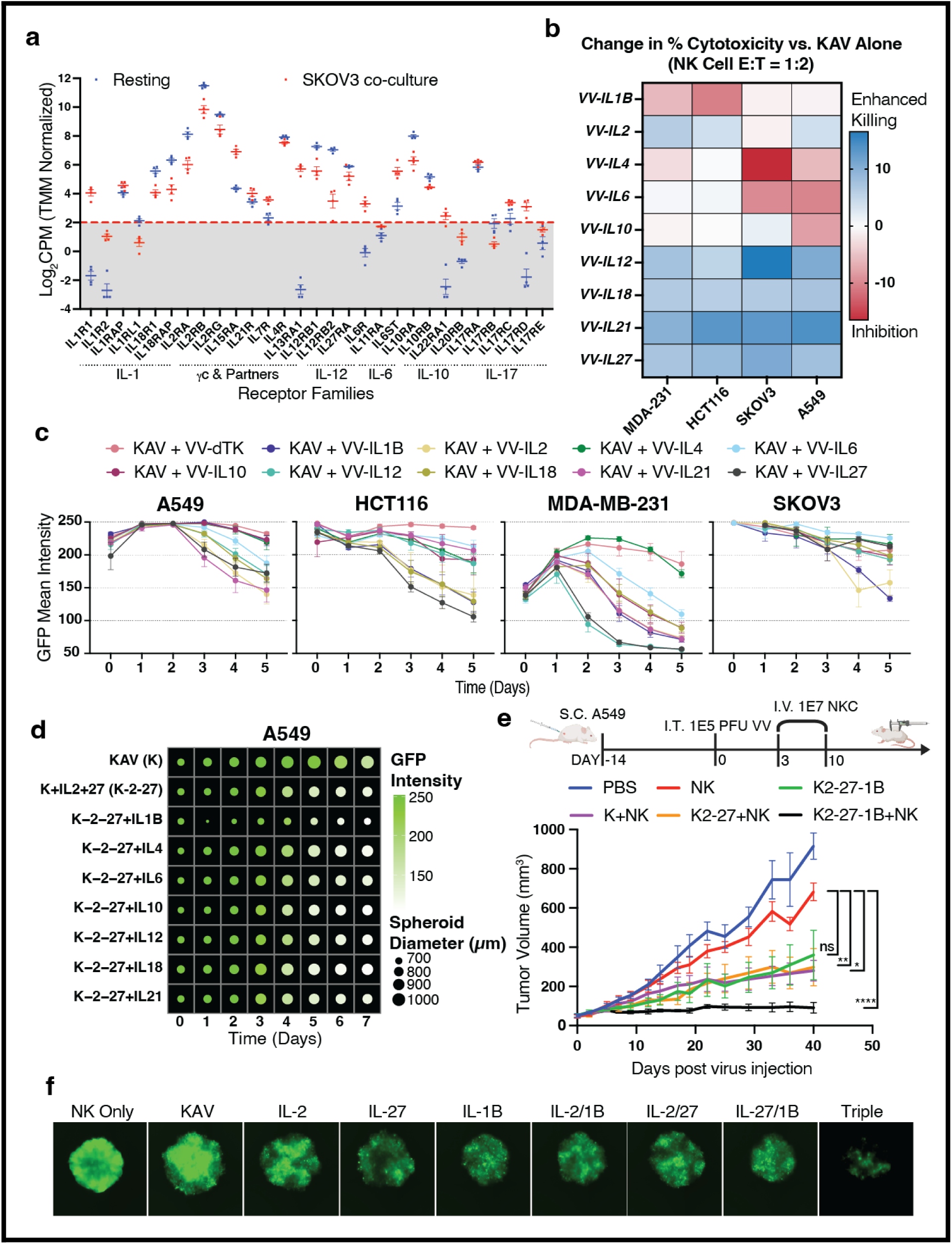
Sequential receptor-guided screening identifies complementary signals that sustain NK-cell function at limiting effector density. (A) Interleukin receptor expression profile in resting (blue) and SKOV-3 co-cultured (red) EFT NK cells. Log2 counts per million (CPM) values (TMM normalized) are shown for receptors categorized by family: IL-1, yc and partners, IL-12, IL-6, IL-10, and IL-17. Data represent n = 4 independent biological donors per condition, batch-corrected across two independent experiments. The horizontal dashed line indicates a detection threshold of 2 Log2CPM. Data are presented as individual points with mean +/- SEM. (B) Heatmap showing the change in percentage cytotoxicity of EFT NK cells (E:T 1:2) across four cell lines when KAV is combined with the indicated interleukin-expressing viruses. Values represent the mean difference in lysis relative to a KAV plus VV-Ctrl baseline (n = 4 independent biological donors: ND602, ND608, ND657, and ND688). Blue denotes enhanced killing and red denotes inhibition. (C) Longitudinal monitoring of 3D tumor spheroids across four cell lines (A549, HCT116, MDA-MB-231, and SKOV3) treated with KAV in combination with indicated interleukin-expressing viruses. Spheroids were infected (MOI 0.1) and cocultured with IL-2 deprived EFT NK cells at a 1:5 E:T ratio. GFP mean intensity was measured daily over 5 days. Data represent mean +/- SEM (n = 5 independent biological donors: ND502, ND602, ND616, ND657, and ND688). (D) Simulation of A549 3D spheroid growth and viability over 7 days following treatment with KAV or combinatorial cytokine strategies. Spheroids were treated with KAV (K), K-2-27 (KAV + VV-IL-2 + VV-IL-27), or K-2-27 in combination with the indicated interleukin-expressing viruses (IL-X). Dot size represents the average spheroid diameter (um), and color intensity represents mean GFP intensity. The GFP intensity scale is represented from 100 (white) to 250 (green). Values represent the average of n = 4 independent biological donors (ND602, ND616, ND657, and ND669). (E) In vivo evaluation of combinatorial cytokine strategies in an A549 subcutaneous xenograft model. Top: Experimental timeline indicating tumor seeding (day -14), I.T. viral injection (1 x 10^5 PFU total, day 0), and I.V. EFT NK cell injections (1 x 10^7 cells per dose) on days 3 and 10 post-infection. Bottom: Tumor volume measurements followed through day 40 post-virus injection. Cocktail treatments consist of KAV, VV-IL-2, VV-IL-27, and VV-IL-1β (2.5 x 10^4 PFU each). Data represent mean +/- SEM (n = 5 mice per group); P values were determined by two-way ANOVA. (F) Deconvolution of cytokine effects on EFT NK cell function in 3D A549 spheroids at a 1:10 E:T ratio. Representative Celigo fluorescence images showing spheroid GFP expression at day 6 post-treatment. All VV expressing cytokine viruses (IL-2, IL-27, IL-1β, and combinations) were administered in conjunction with KAV. “Triple” denotes the combination of KAV with VV-IL-2, VV-IL-27, and VV-IL-1β. Images are representative of one independent biological donor.

We next asked whether the signaling program selected through this sequential screening strategy retained activity in vivo. In a subcutaneous A549 xenograft model, multiple KAV and VV-Cyt combinations with NK cells improved tumor control relative to NK cells alone, whereas the complete KAV, IL-2, IL-27, and IL-1β combination produced the greatest reduction in tumor burden (Fig. 4E and fig. S10). Finally, deconvolution at a 1:10 E:T ratio further illustrated the enhanced activity of the combined signals prioritized through the sequential screen (Fig. 4F).

### Selected cytokine modules provide complementary cytotoxic and metabolic support

To define the functional contributions of each cytokine, we profiled EFT NK cells following exposure to conditioned media from virus-infected tumor cells (Fig. 5A). While the KAV baseline supported NK-cell function, the addition of IL-27 promoted a transcriptional shift toward a mature effector state associated with enhanced cytotoxicity (Fig. 5B and fig. S11A). IL-2 and IL-1β produced distinct transcriptional programs. IL-2 preferentially enriched oxidative phosphorylation-associated pathways, whereas IL-1β induced signatures associated with leukocyte trafficking and cellular stress responses, including upregulation of EPAS1 (HIF-2α). Metabolic assays further showed that inclusion of IL-1β in this conditioned environment reduced maximal oxygen consumption rate (OCR) (Fig. 5C).

**Fig. 5.**
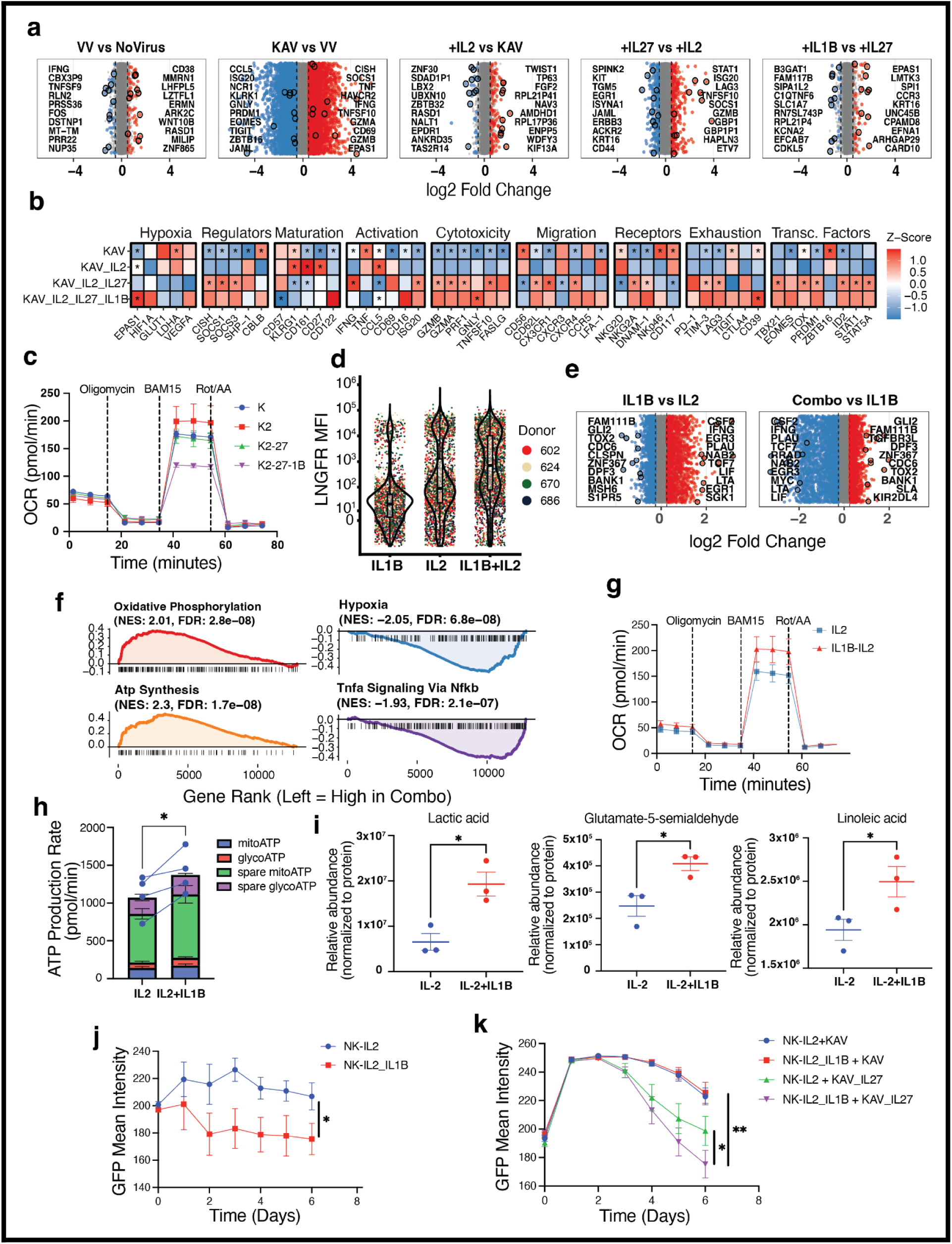
Selected cytokine modules provide complementary cytotoxic and metabolic support to EFT NK cells. (A) Volcano strip plots of differential gene expression in EFT NK cells after 48 h incubation with filtered supernatants from infected A549 cells (72 h post-infection, MOI 0.05). Points represent genes meeting significance thresholds from a prioritized functional list. (B) Heatmap of gene Z-scores organized by functional modules. (C) Oxygen consumption rate (OCR) of conditioned EFT NK cells measured at baseline and following sequential treatment with oligomycin A, BAM15, and rotenone/antimycin A. (D) Cell-by-cell quantification of CD271 (tLNGFR) surface expression on EFT NK cells following retroviral transduction on day 5 and expansion. (E) Volcano strip plots comparing the transcriptomes of CD271-selected NK cells engineered to express IL-1β, IL-2, or the Combo (IL-1β and IL-2). (F) Gene set enrichment analysis (GSEA) for oxidative phosphorylation, hypoxia, and inflammation-related pathways in Combo versus IL-2 engineered NK cells. (G and H) Metabolic profiling of engineered NK cells showing (G) OCR measurements and (H) calculated mitochondrial and glycolytic ATP production rates derived from the Seahorse Mito Stress Test. (I) Normalized relative abundance of intracellular lactic acid, glutamate-5-semialdehyde, and linoleic acid in IL-2 and Combo (IL-2 + IL-1β) engineered NK cells, measured by LC-HRMS. Each dot represents an individual healthy donor (n = 3); bars show mean ± SEM. *p < 0.05, ratio paired t-test. (J and K) Longitudinal GFP mean intensity of A549 tumor spheroids treated with engineered NK cells at effector-to-target (E:T) ratios of (J) 1:3 without virus or (K) 1:12 in combination with indicated viruses. Unless otherwise specified, data represent mean +/- SEM from n = 4 independent biological donors (ND502, ND607, ND657, ND670 for A–C; 602, 624, 670, 686 for D–J). Asterisks (*) denote P < 0.05 compared to the preceding sequential treatment in (B) or the indicated baseline comparison in (I). Significance thresholds for volcano plots (A, E) were defined as P < 0.05 and |log2 FC| > 0.5.

To further examine the intrinsic effects of IL-2 and IL-1β independently of the viral environment, we engineered EFT NK cells by retroviral transduction on day 5 of production (Fig. 5D). Following expansion, IL-1β-expressing cells were depleted relative to the IL-2 group, whereas cells expressing the combination (Combo) were enriched (Fig. 5D). Transcriptomic profiling showed that IL-1β alone induced a strong inflammatory program that was attenuated by co-expression of IL-2 (Fig. 5E). Comparison of Combo versus IL-2-expressing cells (fig. S11B) identified enrichment of oxidative phosphorylation and ATP synthesis pathways together with reduced hypoxia-and TNFα-associated signaling (Fig. 5F). Consistent with these transcriptional changes, Seahorse assays showed increased maximal OCR and total ATP production in the Combo group (Fig. 5, G and H).

Semi-targeted metabolomic profiling further identified significant elevations in lactic acid, glutamate-5-semialdehyde, and linoleic acid in the Combo cohort (Fig. 5I). Increased lactate is consistent with enhanced glycolytic activity (*39*), while elevated linoleic acid has been associated with mitochondrial lipid metabolism (*40*). Glutamate-5-semialdehyde, an intermediate linked to glutamine-derived proline and ornithine metabolism, further suggested broader metabolic remodeling. This pattern was accompanied by a trend toward increased branched-chain amino acids, including leucine and valine (*41, 42*) (fig. S12). Functionally, Combo-engineered NK cells demonstrated improved tumor control in 3D spheroids and retained activity at limiting E:T ratios when combined with KAV and VV-IL27 (Fig. 5, J and K). Collectively, these findings support complementary effects of IL-2 and IL-1β on the metabolic and functional state of EFT NK cells and provide a rationale for their combined inclusion in the engineered viral platform.

### Consolidation of selected signals into a single viral chassis reveals a potency–toxicity tradeoff

To consolidate the multi-cytokine strategy into a single delivery system, we generated a second-generation virus (KAV2) by inserting IL-1β, IL-2, and IL-27 into the I4L locus of the KAV backbone (Fig. 6A). To establish a genetically matched control for I4L disruption, the original single-locus platform, encoding CXCL11 and sushi-IL-15Rα/IL-15 in the J2R locus, was constructed with an equivalent I4L deletion and designated KAV1. Western blot and ELISA confirmed preserved multiplex cytokine expression by KAV2, with each transgene expressed at levels comparable to its respective single-cytokine control virus (fig. S13, A and B). Multi-step growth kinetics showed that although KAV1 achieved higher titers, KAV2 retained replication kinetics similar to the multi-virus cocktail (Fig. 6B). Functional assessment in 2D and 3D tumor models demonstrated that KAV2 recapitulated the activity of the cytokine cocktail and produced greater antitumor activity than KAV1 or NK-cell monotherapy (Fig. 6C and fig. S13C). KAV2 also enhanced the killing activity of freshly isolated, non-expanded primary NK cells at a 1:2 E:T ratio (Fig. 6D).

**Fig. 6.**
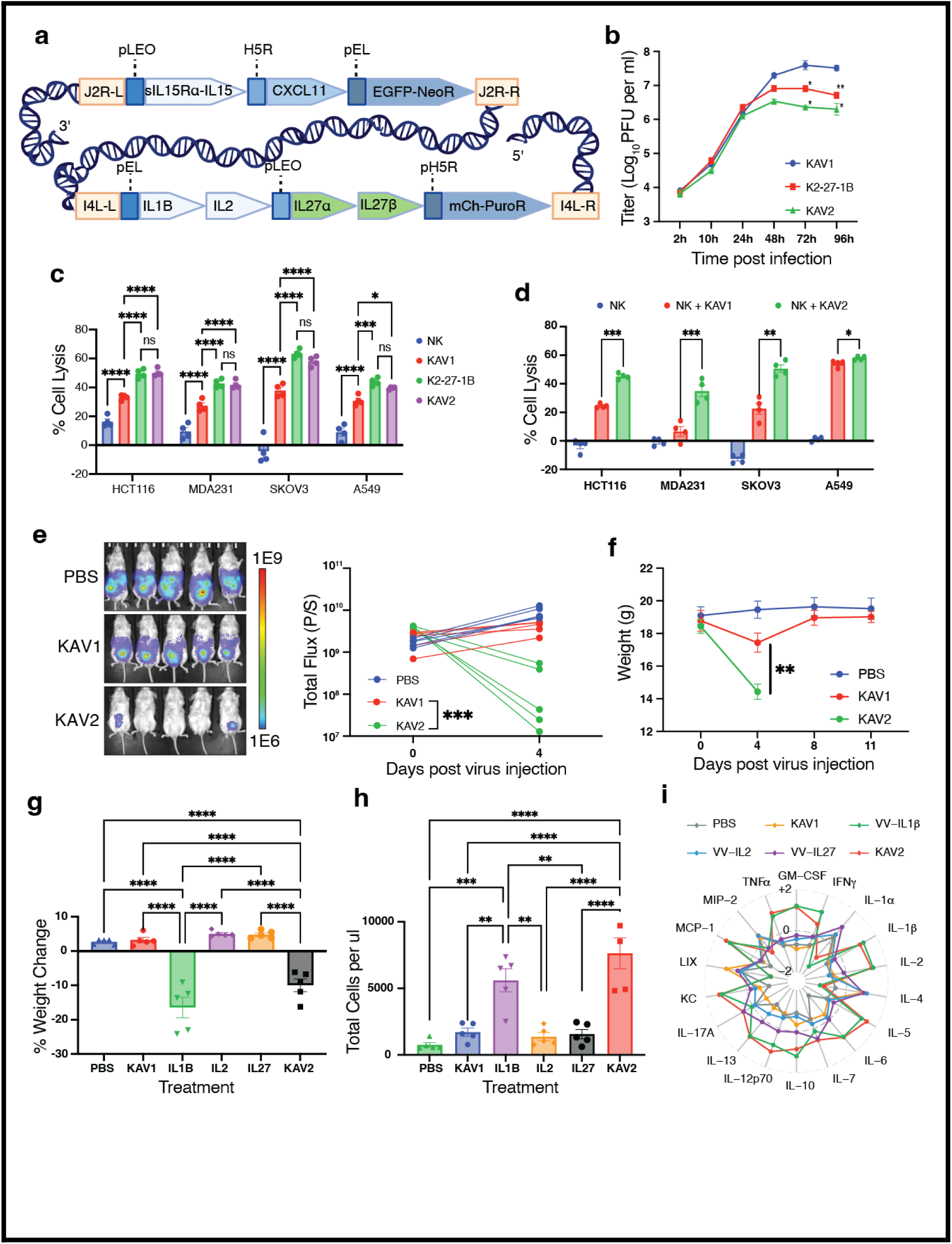
Combining multiple cytokines into a single oncolytic virus enhances anti-tumor activity but triggers systemic toxicity. (A) Schematic of the KAV2 platform design. KAV2 was generated by integrating IL-1β, IL-2, and IL-27 into the I4L locus of the KAV1 backbone (which contains CXCL11 and sIL15Ra-IL15 in the J2R locus). (B) In vitro viral fitness kinetics. A549 cells were infected (MOI 0.01) with KAV1, the four-virus cocktail (K2-27-1B), or KAV2; total viral progeny were harvested at indicated time points and titered on U2OS cells. (C) 2D cytotoxicity of EFT NK cells at an effector-to-target (E:T) ratio of 1:5 against indicated solid tumor lines. (D) 2D cytotoxicity of freshly isolated, non-expanded human NK cells at a 1:2 E:T ratio. (E and F) Evaluation of systemic toxicity in an HCT116 intraperitoneal (I.P.) xenograft model. NSG mice were seeded with 1 x 10^6 HCT116 cells and treated 5 days later with a single I.P. injection of indicated viruses (1 x 10^5 PFU). (E) Representative bioluminescence (IVIS) images and (F) mouse weight change 4 days post-virus injection. (G–I) Characterization of the cytokine-driven toxicity profile in a subcutaneous HCT116 model. Mice were treated with 1 x 10^5 PFU of indicated viruses 14 days post-tumor seeding. (G) Percentage weight change 3 days post-virus injection. (H) Absolute Trucount quantification of murine CD45+ cells in peripheral blood 3 days post-virus injection. (I) Radar plot showing the comparative signature of 18 murine cytokines in peripheral blood. Values represent mean Z-scores derived from Log10-transformed Luminex assay data. Unless otherwise specified, in vitro data represent mean +/- SEM from n = 4 independent biological donors. In vivo data represent mean +/- SEM from n = 5 mice per group. Asterisks (*) denote P < 0.05. Statistical significance for multi-group comparisons was determined by one-way or two-way ANOVA with Tukey’s post-hoc test.

Consolidation of these selected signals into a single viral chassis, however, revealed a previously unrecognized safety liability. In an HCT116 intraperitoneal model, KAV2 treatment in the absence of NK-cell administration caused rapid loss of bioluminescent tumor signal together with severe weight loss (Fig. 6, E and F). While three of the four human cytokines evaluated in this study (excluding IL-27) are known to at least partially cross-activate their corresponding murine receptors (*43–45*), human IL-1β is equipotent in mice (*43, 46*) and is a potent mediator of systemic inflammation, fever, and acute-phase responses in both species (*47*). Consistent with this cross-species activity, comparison of the individual cytokine-expressing viruses in a subcutaneous HCT116 model showed that VV-IL-1β induced weight loss comparable to KAV2, whereas the other cytokine-expressing viruses did not (Fig. 6G). This phenotype was accompanied by a marked increase in circulating murine CD45+ cells and a systemic cytokine profile closely resembling that induced by KAV2 (Fig. 6, H and I and fig. S14). Together, these findings identify constitutive IL-1β expression as the major inflammatory liability of the consolidated platform and establish a need for temporal control of the high-potency cytokine module.

### Inducible control of the IL-2/IL-1β module preserves antitumor activity while limiting systemic toxicity

To address the inflammatory liability identified after consolidation of the selected signals into KAV2, we engineered an inducible version of the platform (iKAV). The J2R and I4L modifications were retained, while the IL-1β-and IL-2-linked open reading frame was placed under the control of a doxycycline-inducible promoter together with constitutive expression of the tetracycline repressor (TetR) (Fig. 7A). ELISA analysis confirmed approximately 100-fold induction of IL-1β and IL-2 following doxycycline treatment, whereas IL-27 expression remained unchanged because it was not placed under inducible control (Fig. 7B). In 3D spheroid models across A549, HCT116, and SKOV3 cells, induced iKAV recapitulated the enhanced activity of KAV2 at a limiting 1:10 E:T ratio, whereas uninduced iKAV showed activity comparable to KAV1 (Fig. 7C). These findings demonstrated that the high-potency phenotype could be placed under external temporal control without altering the remaining components of the platform.

**Fig. 7.**
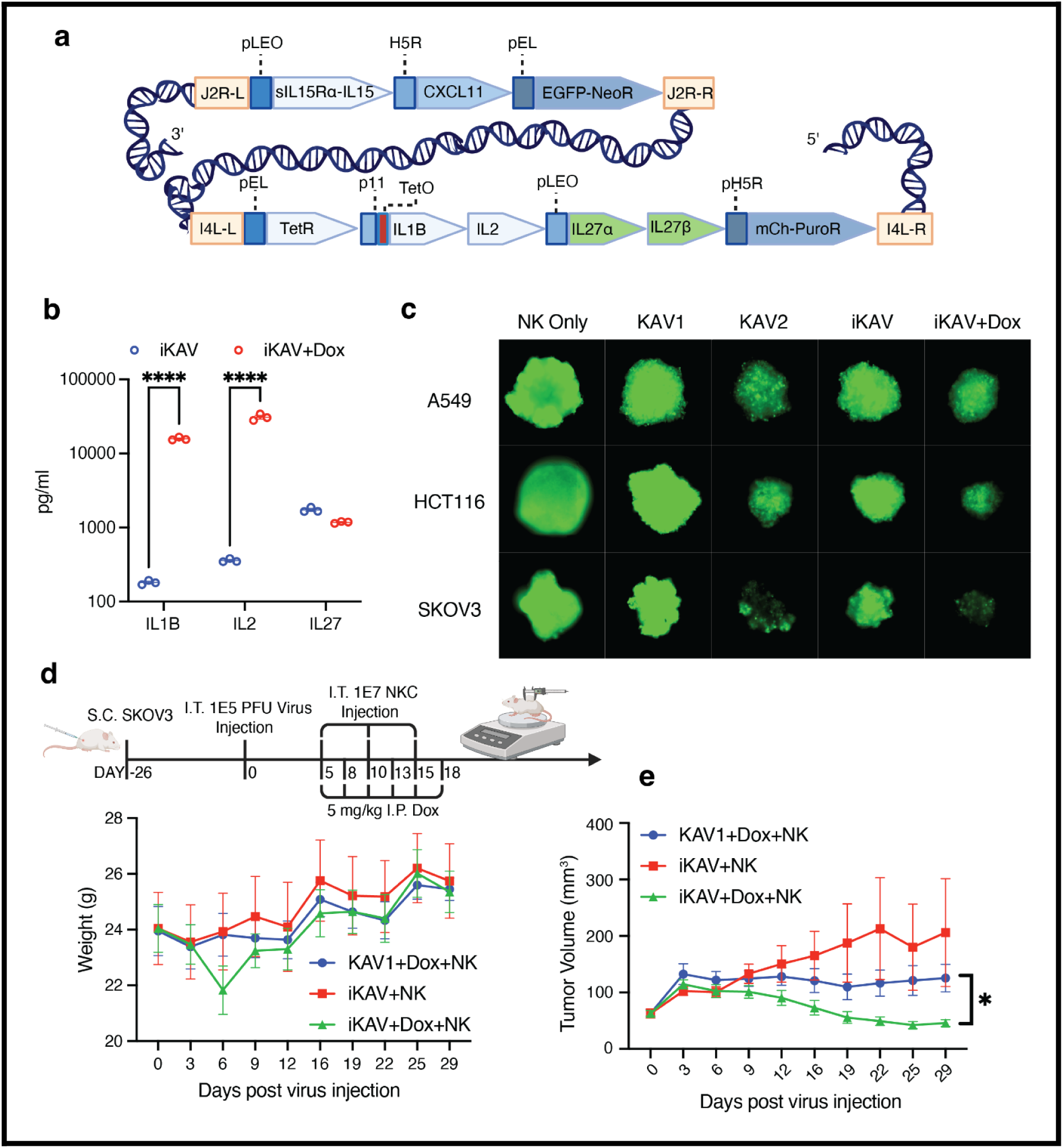
Inducible control of the IL-2/IL-1β module preserves antitumor activity while limiting systemic toxicity. (A) Architectural schematic of the programmable iKAV (inducible Killer Assistant Virus) viral genome. This optimized chassis utilizes dual deletions within the thymidine kinase (*J2R*) and ribonucleotide reductase (*I4L*) loci. The payload, featuring a 2A-linked human IL-2/IL-1β module, is governed by a modified late promoter containing a tetracycline operator (*p11-TetO*). Precise temporal regulation is achieved via constitutive expression of the tetracycline repressor (TetR) under the control of a synthetic early/late (*pEL*) vaccinia promoter. (B) ELISA-based measurement of human IL-1β, IL-2, and IL-27 concentrations in filtered A549 cell supernatants 48 h post-infection (MOI 0.1) with iKAV. Transgene output was evaluated following the addition (+Dox) or withdrawal (-Dox) of 1 μg/mL doxycycline. Data are mean ± SEM from n = 3 replicates. (C) Representative Celigo fluorescence images of A549, HCT-116, and SKOV-3 3D tumor spheroids at 7 days, 7 days, and 4 days post-treatment, respectively. Spheroids expressing EGFP were monitored for tumor viability at a limiting 1:10 E:T ratio. Efficacy is compared across NK cell monotherapy, KAV1, KAV2, and the programmable iKAV platform in both uninduced (–Dox) and induced (+Dox) states. (D) In vivo weight and (E) Average tumor volume in NSG mice bearing established subcutaneous SKOV-3 ovarian tumors. Following treatment with intratumoral injections of the indicated oncolytic viruses (KAV1 or iKAV), mice received therapeutic infusions of 1 x 10^7 EFT NK cells at indicated time points. Experimental cohorts were monitored under conditions of doxycycline withdrawal (-Dox) or intermittent administration (+Dox; 5 mg/kg I.P. on days 5, 8, 10, 13, 15, and 18). Data are mean +/- SEM (n = 6 (KAV1) or 7 (iKAV) mice per group). Asterisks (*) denote P < 0.05. Statistical significance for multi-group comparisons was determined by two-way ANOVA

We next evaluated whether inducible control could retain antitumor activity while reducing the systemic toxicity associated with constitutive cytokine expression. In a subcutaneous SKOV3 xenograft model, intermittent doxycycline induction of iKAV improved tumor control relative to the uninduced platform while maintaining a stable body-weight profile (Fig. 7, D and E). Together, these results show that temporal regulation of the IL-2/IL-1β module retains the functional advantage of the multiplexed platform while limiting the systemic toxicity associated with constitutive expression.

## DISCUSSION

Over the past 15 years, considerable efforts have sought to bridge the systemic potency of adoptive cell therapies with the localized delivery capabilities of oncolytic viruses (OVs) (*48*). These strategies have used the genomic capacity of large DNA viruses, including adenovirus, herpes simplex virus, and vaccinia virus, to support T-or NK-cell therapies through transgenes for tumor chemoattraction (*49–51*), antigen painting (*52–55*), or localized immune activation (*55, 56*). However, OV–cell therapy combinations have generally been designed by adding established immunostimulatory payloads to a selected viral platform rather than by beginning with the biological state and functional requirements of the manufactured cell product. Likewise, viral chassis selection has rarely been evaluated as a function of compatibility with the activated effector population itself (*48*). Here, we instead used the manufactured NK-cell product as the design input: defining its functional liabilities, selecting a viral chassis that retained function and spread under NK-cell pressure, and iteratively engineering localized support signals around those requirements.

Within NK-cell–OV combinations, translational performance is strongly influenced by the cell source and its manufacturing state. Many preclinical studies rely on the NK-92 cell line (*57*); however, the clinical requirement for lethal irradiation restricts its functional activity to a brief, non-replicative window (*58, 59*). In primary NK-cell approaches, strategies to enhance trafficking have included engineering the cells to express exogenous homing receptors, such as CCR5 (*49*), thereby placing the trafficking solution directly within the cell product. An alternative approach is to map the endogenous receptor landscape of the manufactured product and engineer the viral niche to complement it. Crucially, preclinical studies have largely bypassed cryopreserved effectors, despite their central role in scalable, off-the-shelf therapies (*8*). Because expansion and cryopreservation can reshape trafficking-related receptor expression and functional fitness, microenvironmental support should be designed around the post-thaw state of the therapeutic product rather than an idealized, freshly isolated effector cell.

To address these hurdles, we focused on an off-the-shelf product of expanded, frozen, and thawed (EFT) NK cells preconditioned with high-dose IL-15 and IL-18 (*15*). While this manufacturing regimen prevents post-thaw fratricide and preserves high viability, prolonged ex vivo exposure to supraphysiologic cytokines creates an acute dependency: upon thaw, these effectors lose sustained killing capacity at limiting effector-to-target ratios and require continuous extrinsic cytokine support. Pairing such an activated innate effector with an oncolytic virus introduces a fundamental biological constraint: the viral platform must withstand the same antiviral machinery that gives the cell product its therapeutic function. In direct head-to-head evaluation, HSV-1 was rapidly restricted under EFT NK-cell pressure, whereas oncolytic vaccinia virus (VV) retained replication and spread, establishing a resilient chassis around which to construct an extrinsic support niche.

With oncolytic vaccinia validated as a compatible chassis, we next leveraged the phenotypic profile of EFT NK cells to direct the engineering of the initial support module. Consistent, high-level expression of CXCR3 motivated the incorporation of its cognate chemokine, CXCL11, to complement the manufacturing-associated reduction in chemokine responsiveness (*21*), while a sushi-IL-15Rα/IL-15 complex was integrated to provide survival and homeostatic support. This first-generation killer assistant virus (KAV1) significantly enhanced EFT NK-cell activity, but its benefit diminished at limiting effector-to-target ratios. Moreover, attempting to overcome this threshold by simply escalating the adoptive cell dose encountered a clear safety ceiling, as high cellular doses triggered systemic toxicity in vivo. These findings showed that localized trafficking and survival cues established an important baseline but were insufficient to sustain effector function under high tumor burden, creating the need for additional complementary support signals.

Because the baseline protection of KAV1 diminished at sparse effector densities, we used the endogenous interleukin receptor repertoire of EFT NK cells to constrain candidate payloads for secondary engineering. Rather than evaluating arbitrary cytokine combinations, we subjected this receptor-informed viral library to sequential functional screening in three-dimensional tumor spheroids under escalating selective stringency. This screening hierarchy first prioritized IL-2 and IL-27 at a 1:5 effector-to-target ratio; increasing the stringency to 1:10 subsequently revealed IL-1β as the strongest additional signal when added to the KAV1/IL-2/IL-27 baseline. These factors emerged through iterative functional selection under limiting effector conditions rather than empirical guesswork, and their combined activity was recapitulated in vivo using a multi-virus cocktail. These findings show that stepwise increases in selective pressure can expose support requirements that remain less apparent under more permissive assay conditions.

Downstream profiling of the prioritized signals revealed distinct and complementary functional programs. While IL-15 established baseline homeostatic and survival support for post-thaw NK-cell function, IL-27 was associated with a stronger cytotoxic effector program. This distinction is consistent with prior work separating IL-27 signaling from more strongly inflammatory drivers such as IL-12, which can accelerate activation-induced cell death (*60*). In contrast, the IL-2/IL-1β module produced more pronounced changes in metabolic and stress-associated programs than in canonical cytotoxicity markers. The bioenergetic impact of IL-1β was context-dependent: exposure within tumor-conditioned media reduced maximal respiration, whereas co-expression with IL-2 increased maximal oxygen consumption, elevated total ATP production, and altered the abundance of several metabolic intermediates. Together, these findings indicate that the selected factors engage complementary cytotoxic and bioenergetic programs, providing a biological rationale for their consolidation into a unified viral support niche.

Consolidating these selected signals into a single viral chassis exposed an intrinsic bioengineering challenge: payload integration coupled therapeutic potency with systemic toxicity. While KAV2 preserved the antitumor activity of the multi-virus cocktail, constitutive expression of human IL-1β triggered severe systemic inflammation. Human IL-1β is known to retain biological activity in mice (*46*), and its continued production from the replicating viral platform was associated with the systemic inflammatory liability observed in vivo. Rather than discarding a factor identified through stringent functional selection, we addressed this exposure constraint by placing the IL-2/IL-1β module under doxycycline-inducible control. This shift from static payload delivery to conditional regulation illustrates a broader design principle for multiplexed cell-therapy support: as complementary signals are consolidated into increasingly potent therapeutic systems, emergent safety liabilities may need to be managed at the level of payload control rather than by further altering the selected biology.

Evaluating this platform in NSG xenograft models provided a controlled setting to interrogate the interactions among human tumor targets, human EFT NK cells, and the engineered viral niche without host-mediated rejection of the human components. However, this model necessarily excludes endogenous immune compartments, including myeloid and adaptive populations, that influence viral clearance and cellular immunotherapy in intact hosts. Furthermore, species-specific differences in NK-cell biology (*61*) and variable cross-reactivity of human cytokines in mice constrain direct physiological extrapolation. While lymphodepletion regimens routinely administered before clinical adoptive cell transfer transiently reduce endogenous immune pressure (*62*), future studies in humanized models or appropriately matched immunocompetent surrogate systems will be important for defining how this localized oncolytic niche interacts with a complete host immune environment.

Collectively, these findings establish an engineering framework that uses the phenotypic and functional state of a manufactured cell product to direct the rational construction of a complementary oncolytic niche. Rather than placing every supportive function within the cellular product itself, liabilities in trafficking, survival, cytotoxicity, and metabolic fitness can be addressed through signals delivered locally from a programmable viral platform. In this work, that framework integrated cell-product profiling and viral chassis selection with receptor-informed payload prioritization, functional screening under limiting effector conditions, and inducible genetic control of an emergent toxicity. More broadly, this paradigm provides a blueprint for co-engineering adoptive cell therapies and oncolytic platforms as interdependent components of a unified therapeutic system, with the extrinsic microenvironment tailored to the biological requirements of the manufactured cell product.

## MATERIALS AND METHODS

### Study Design

The primary objective of this study was the iterative development of an OV engineered to provide a localized, multi-cytokine signaling niche capable of sustaining the effector function of cryopreserved, ex vivo–expanded human NK (EFT NK) cells within solid tumor environments. Primary human NK cells were isolated from healthy donor peripheral blood mononuclear cells (PBMCs) obtained through the University of Pennsylvania Center for Cellular Immunotherapies (CCI) Human Immunology Core (HIC). Purification was achieved via negative magnetic selection using a standardized enrichment kit (Miltenyi Biotec). For in vitro functional characterization, 3 to 5 independent biological replicates were utilized; each replicate consisted of NK cells from a distinct healthy donor, which were expanded and cryopreserved at separate intervals to ensure experimental robustness and cross-donor consistency during simultaneous testing. In vivo therapeutic efficacy was evaluated using murine xenograft models with a sample size of n = 5 mice per treatment group. Investigators were not blinded to the treatment assignments during the course of the study.

### Cell Lines and Viruses

Human solid tumor cell lines SKOV3, A549, HCT116, and MDA-MB-231 were obtained from the ATCC and engineered to express click beetle green luciferase (CBG) for bioluminescence-based cytotoxicity assays. The identity of all cell lines was verified by STR profiling (ATCC), and they were confirmed negative for mycoplasma (Cambrex MycoAlert) following culture without antibiotics. Cells were cultured in RPMI 1640 (Thermo Fisher) or DMEM Glutamax supplemented with 10% fetal bovine serum (FBS) (Seradigm) and penicillin-streptomycin (Gibco).

Wild-type vaccinia virus (VV; Lister strain) and herpes simplex virus-1 (HSV-1; KOS strain) were obtained from the ATCC and used as backbones for subsequent modification. Recombinant viruses were generated using a high-throughput, antibiotic-driven parallel engineering platform as previously described (*63*). Homologous recombination was performed in HEK293 cells using donor plasmids containing the transgene and a selection-marker cassette (PuroR-T2A-EGFP or NeoR-T2A-mCherry). Recombinant clones were enriched through sequential passaging and 1-2 rounds of plaque purification in U2OS cells under metabolic selection with Puromycin or Neomycin. Transgenes were targeted to the I4L or TK (J2R) loci in VV and the UL39 (ICP6) locus in HSV-1. For head-to-head comparison, virus titers were determined on Vero cells; for the remainder of the study, VV was titered on HeLa or U2OS cells. High-titer stocks for in vivo studies were produced in Vero cells (HSV-1) or HeLa cells (VV) and purified via sucrose cushion centrifugation.

### NK Cell Purification, Expansion, and Cryopreservation

Primary human NK cells were isolated from healthy donor PBMCs and purified via negative magnetic selection (Miltenyi Biotec). For expansion, NK cells were co-cultured at a starting 1:10 ratio with irradiated (100 Gy) K562 feeder cells expressing 4-1BBL, CD40, and membrane-bound IL-21 (K-IL-21), as previously described (*35*). Cells were maintained in X-VIVO 10 (Lonza) supplemented with 10% human AB serum and 100 U/mL IL-2 in G-Rex 6-well or 24-well plates (Wilson Wolf Manufacturing). High-dose IL-2 expanded (2E) control cells were cultured similarly but in the presence of 1000 U/mL IL-2 without feeder support. The expanded-frozen-thawed (EFT) product was generated by treating expanded cells on day 15 with 50 ng/mL IL-15 and 250 ng/mL IL-18 (InvivoGen) overnight prior to cryopreservation in CryoStor CS10. Cryopreserved cells were thawed and rested overnight in X-VIVO 10 with 10% human AB serum and 100 U/mL IL-2 before functional evaluation. For retroviral engineering, BaEV-pseudotyped retroviral particles were produced using 293Vec-BaEV packaging cells (Biovec Pharma). NK cells were transduced on day 5 of expansion using a Retronectin-coated (Takara Bio) 24-well plate protocol; virus was pre-loaded by centrifugation followed by the addition of 0.5 × 10⁶ NK cells per well. Transduced cells were transferred back to G-Rex plates 24 hours post-infection for continued expansion.

### In Vitro Functional assays

For 2D cytotoxicity assays, co-cultures were performed in 24-well plates and subsequently transferred to white 96-well assay plates for bioluminescence measurement. D-luciferin was added at a final concentration of 0.6 mg/mL, and luminescence from CBG-expressing tumor cells was recorded using a plate reader with two to three technical replicates per experimental condition.

Tumor spheroids were generated by seeding 5,000 EGFP expressing cells per well in Biofloat 96-well U-bottom plates (Sarstedt). The next day, spheroids were infected with indicated recombinant viruses, and EFT NK cells were added 4 hours post-infection. Tumor viability was monitored by measuring GFP intensity using a Celigo Image Cytometer (Revvity).

NK cell chemotaxis was evaluated using 5 micrometer pore Transwell inserts (Corning). EFT NK cells were stained with DiD lipophilic dye (Thermo Fisher) and placed in the upper chamber, while conditioned media from virus-infected cells was added to the lower chamber. The number of migrated cells was quantified after a 1-hour incubation period.

### Metabolic Profiling

Mitochondrial function was assessed using a Seahorse XFe96 analyzer (Agilent) with the XF T Cell Metabolic Profiling Kit. EFT NK cells (1.5 x 10^5 cells/well) were seeded in Seahorse XF RPMI Medium supplemented with 10 mM glucose, 2 mM L-glutamine, and 1 mM pyruvate, and OCR and ECAR were measured at baseline and following sequential injection of oligomycin A (1.5 uM), BAM15 (2.5 µM), and rotenone/antimycin A (0.5 µM). For metabolomic profiling, cell pellets were rinsed in ice-cold 0.9% NaCl and extracted by sonication in an acetonitrile/methanol/water (2:2:1) solution with 0.1% formic acid, followed by the addition of 15% NH4HCO3. Protein and cell debris were precipitated at 21,000 rcf. Supernatants were dried under nitrogen, resuspended in acetonitrile/water (1:1), and analyzed on a ZIC-pHILIC column using a Vanquish Duo UHPLC coupled to a Q Exactive Plus mass spectrometer (Thermo Fisher Scientific). Data were acquired in polarity-switching mode (70–1000 m/z) and processed using Tracefinder software with a 5 ppm mass tolerance window.

### In Vivo Models

All animal studies were performed under an approved Institutional Animal Care and Use Committee (IACUC) protocol at the University of Pennsylvania (Philadelphia, PA). NOD/scid/IL-2rg-/-(NSG) mice were originally obtained from The Jackson Laboratories and were maintained by the Stem Cell and Xenograft Core at the University of Pennsylvania in pathogen-free conditions. Eight to 10 weeks old male or female mice were used for all studies.

Tumor growth was monitored twice weekly by bioluminescence imaging (BLI) using the IVIS Lumina III In Vivo Imaging System (PerkinElmer). Recombinant viruses were administered via intratumoral (IT) or intraperitoneal (IP) injection. EFT NK cells were administered via IT or IV injection following viral infection. For inducible KAV (iKAV) experiments, doxycycline was administered via IP injection to trigger transgene expression. For BLI, mice were anesthetized and received an IP injection of D-luciferin (150 mg/kg). Images were captured 5 minutes post-injection and analyzed using AuraAnalysis software. Total tumor flux (photons per second) was used to compare tumor progression across groups.

### Flow Cytometry and Phenotyping

Flow cytometric data were acquired using a BD Symphony A3 Lite (BD Biosciences). A comprehensive list of all antibodies is provided in Supplementary Table 1. Flow data analysis was performed using FlowJo v10. For the cell-by-cell quantification of LNGFR expression shown in Figure 5, compensated data were exported from FlowJo and processed in R using the flowCore and tidyverse packages for data extraction, tidying, and visualization.

### Transcriptomic and Bioinformatic Analysis

Total RNA was sequenced on the Illumina NovaSeq platform. For data presented in Figures 1 through 4, sequencing was performed by Azenta Genewiz, and raw reads were mapped to the human transcriptome using Kallisto. For Figure 5, pre-processed counts-per-million (CPM) data were provided by Plasmidsaurus. All downstream bioinformatic analyses were conducted within the R environment using the Tidyverse framework. Differential expression analysis was performed using the EdgeR and Limma packages. Gene Set Enrichment Analysis (GSEA) and Gene Ontology (GO) terms were evaluated using the fgsea and msigdbr packages, referencing the MSigDB Hallmark gene sets and standard GO categories.

### Statistical Analysis

Statistical analyses were conducted using GraphPad Prism 10 and R, with data generally presented as mean +/-SEM from three to five independent biological donors. Longitudinal tumor growth and in vitro kinetics were evaluated by two-way ANOVA followed by Tukey’s or Sidak’s post-hoc tests. Single-parameter endpoints were analyzed via one-way ANOVA or two-tailed unpaired t-tests, while survival differences were assessed using the log-rank (Mantel-Cox) test. Cytotoxicity EC50 values were calculated using a four-parameter non-linear regression model. For transcriptomic data, differential expression was defined by an adjusted P-value < 0.05 and an absolute log2 fold change > 1. Statistical significance was defined as P < 0.05 across all assays.

## Funding

This work was supported by private funds from the University of Pennsylvania and a ScaleReady G-Rex® Grant. Additional research support was provided by the National Institutes of Health (grant R01-CA278837 to ROC), the Ludwig Research Foundation Princeton Branch (ROC), and the Parker Institute for Cancer Immunotherapy (ROC).

## Author contributions

We would like to thank the Human Immunology Core (HIC) at the Perelman School of Medicine at the University of Pennsylvania, for the provision of PBMCs. The HIC is supported in part by NIH P30 AI045008 and P30 CA016520. We gratefully acknowledge Omar Johnson, Siddhant Kadu, Abdulla Berjis, and Schyler Rowland for their helpful discussions and technical advice concerning the optimization of experimental protocols.

## Competing interests

NCS holds equity in and is a Scientific Advisory Board (SAB) member for BlueWhale Bio, CART Tx, and Waypoint Bio, a SAB member for PanCancer T, and an advisor for Able Sciences, Laverock Therapeutics, and UroGen Pharma. He holds IP related to NK cell cryopreservation. ROC has equity in Nucleus Biologics and Stoic Bio outside of the submitted work, and has patents related to CAR T cell therapy.

## Declaration of Generative AI and AI-assisted technologies in the manuscript preparation process

During the preparation of this work, the authors used ChatGPT and Google Gemini to assist with language editing, improving readability, and polishing manuscript phrasing. The authors critically reviewed, revised, and edited all outputs and take full responsibility for the final content of the published article.

---

**Fig. S1.**
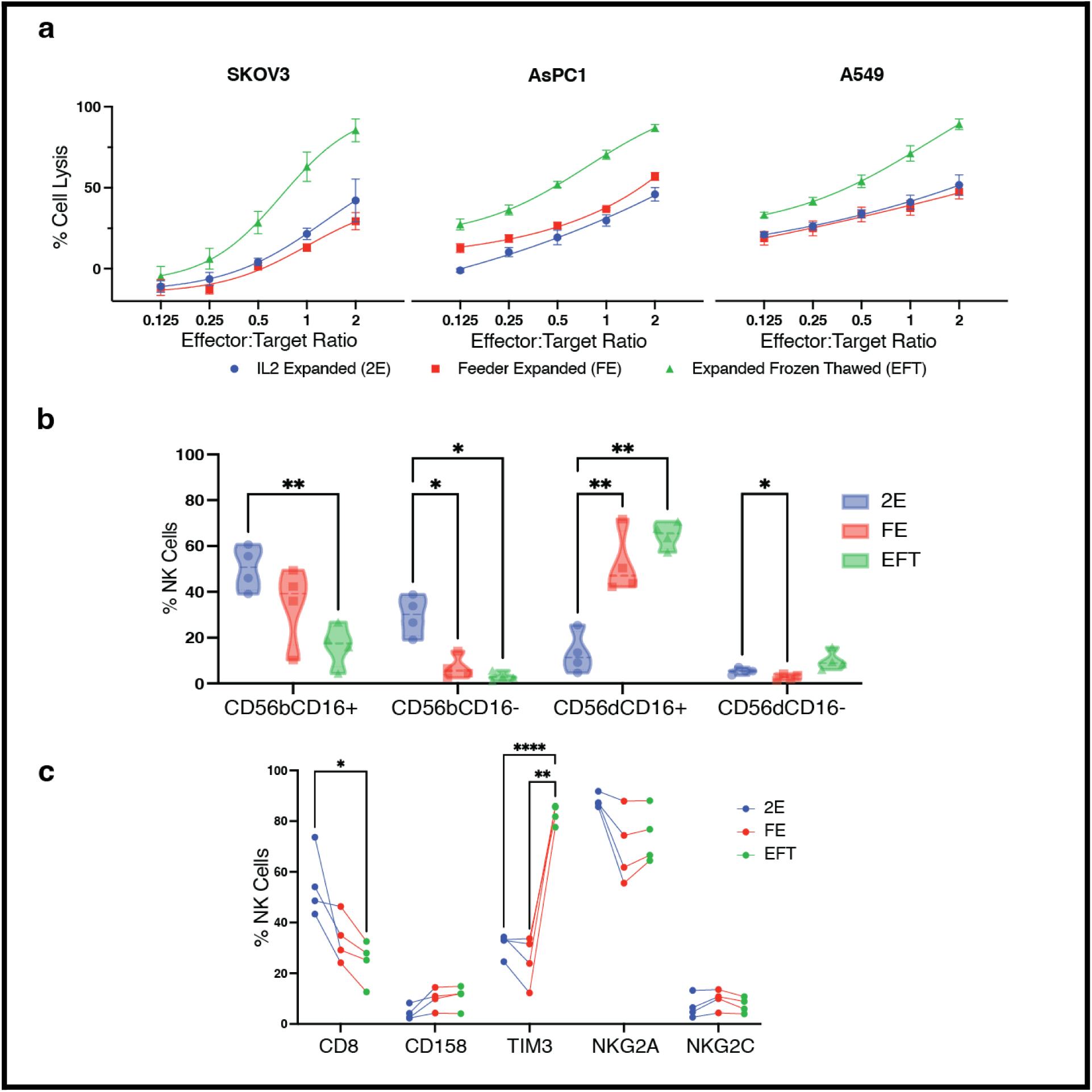
Characterization of EFT NK cell cytotoxicity and phenotypic profile. (A) Cytotoxic activity of IL-2 expanded (2E), feeder-expanded (FE), and expanded frozen-thawed (EFT) NK cells against SKOV3, AsPC1, and A549 cell lines at indicated E:T ratios. Percentage of cell lysis was determined after 24 h. (B) Flow cytometric quantification of NK cell subsets based on CD56 (bright/dim) and CD16 surface expression in 2E, FE, and EFT populations. (C) Surface expression of NK cell markers, including CD8, CD158, TIM3, NKG2A, and NKG2C, across 2E, FE, and EFT populations. Data represent mean +/- SEM (n = 4 donors); P values were determined by two-way ANOVA. *P < 0.05, **P < 0.01, ****P < 0.0001.

**Fig. S2.**
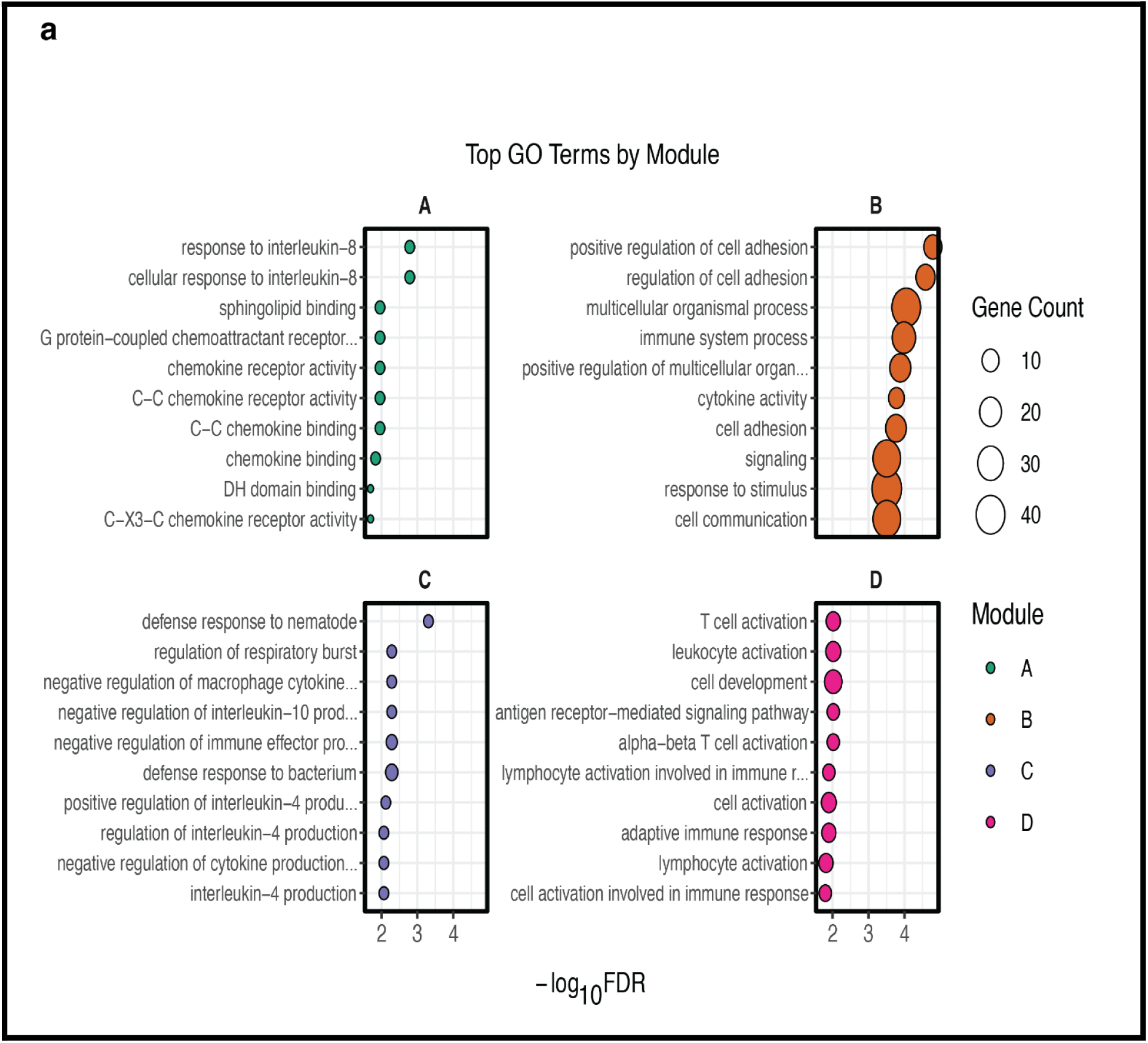
Gene Ontology (GO) enrichment analysis of transcriptomic modules. (A) Enriched GO terms for Module A, highlighting pathways for chemokine receptor activity and response to interleukin-8. (B) Enriched GO terms for Module B, including biological processes for cytokine activity and regulation of cell adhesion. (C) Enriched GO terms for Module C, focusing on defense responses to bacteria and regulation of cytokine production (e.g., IL-4 and IL-10). (D) Enriched GO terms for Module D, centered on pathways for leukocyte and T cell activation. Across all panels, the x-axis represents adjusted P value < 0.01, dot size indicates the gene count associated with each term, and colors designate the corresponding modules A–D.

**Fig. S3.**
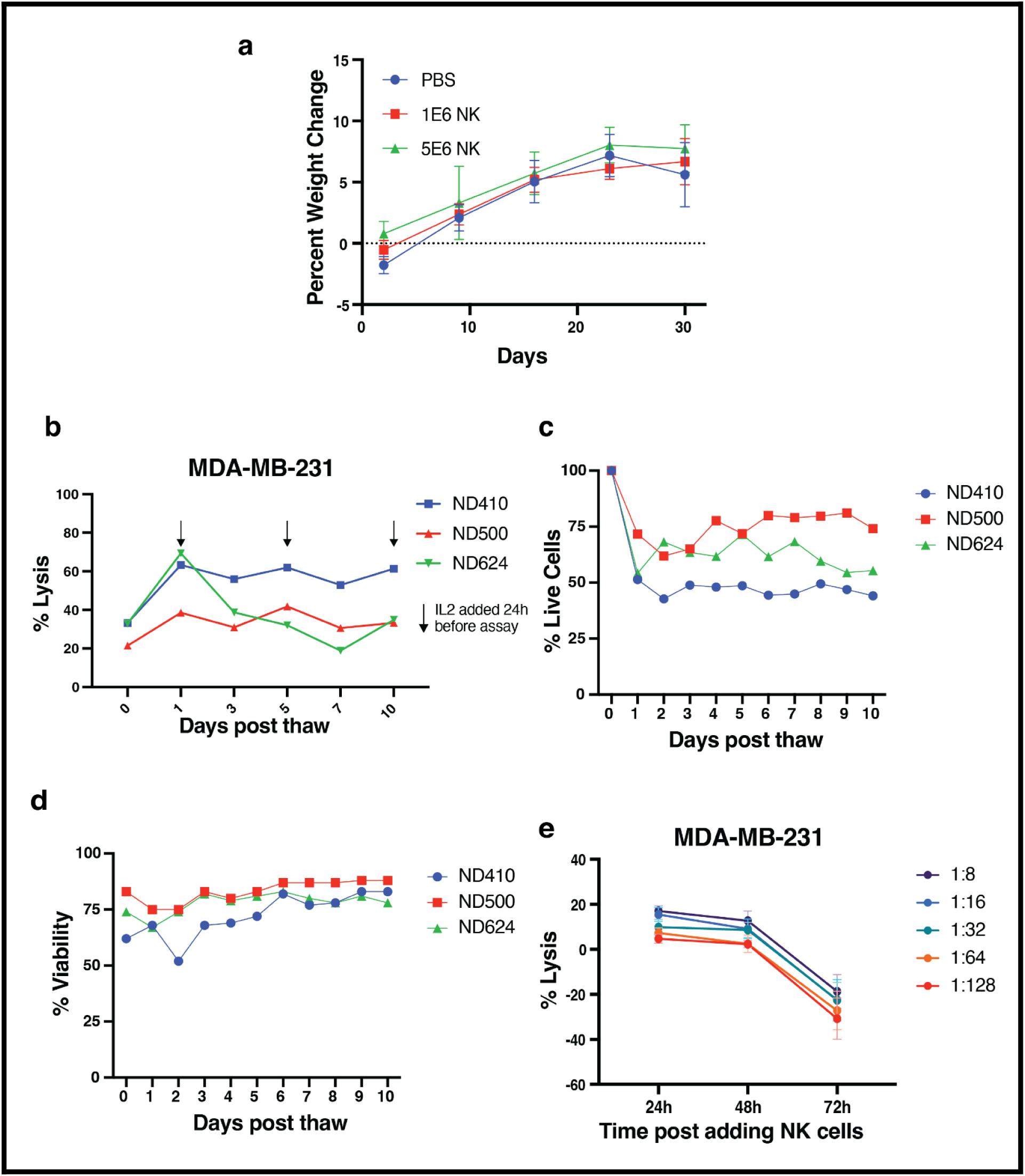
Longitudinal stability, functional kinetics, and in vivo safety of EFT NK cells. (A) Percentage weight change in mice relative to Day 0 following intratumoral injection of PBS, 1 x 10^6 NK cells, or 5 x 10^6 NK cells. Weight was monitored over 30 days. Data represent mean +/- SEM (n = 5 mice per group). (B) Percentage lysis of MDA-MB-231 cells by EFT NK cells at a 1:2 E:T ratio measured over 10 days post-thaw. Arrows indicate time points where IL-2 was added to the media 24 h prior to the assay. Data represent n = 3 independent biological donors (ND410, ND500, and ND624). (C) Percentage of live cells relative to the initial count at Day 0 monitored over 10 days post-thaw. Data represent n = 3 independent biological donors (ND410, ND500, and ND624). (D) Percentage viability of EFT NK cells monitored over 10 days post-thaw as determined by automated cell counting. Data represent n = 3 independent biological donors (ND410, ND500, and ND624). (E) Kinetic analysis of MDA-MB-231 cell lysis at indicated E:T ratios measured at 24 h, 48 h, and 72 h post-coculture. Negative lysis values represent time points where target cell signals exceeded untreated controls. Data represent mean +/- SEM (n = 4 independent biological donors: ND410, ND500, ND607, and ND608).

**Fig. S4.**
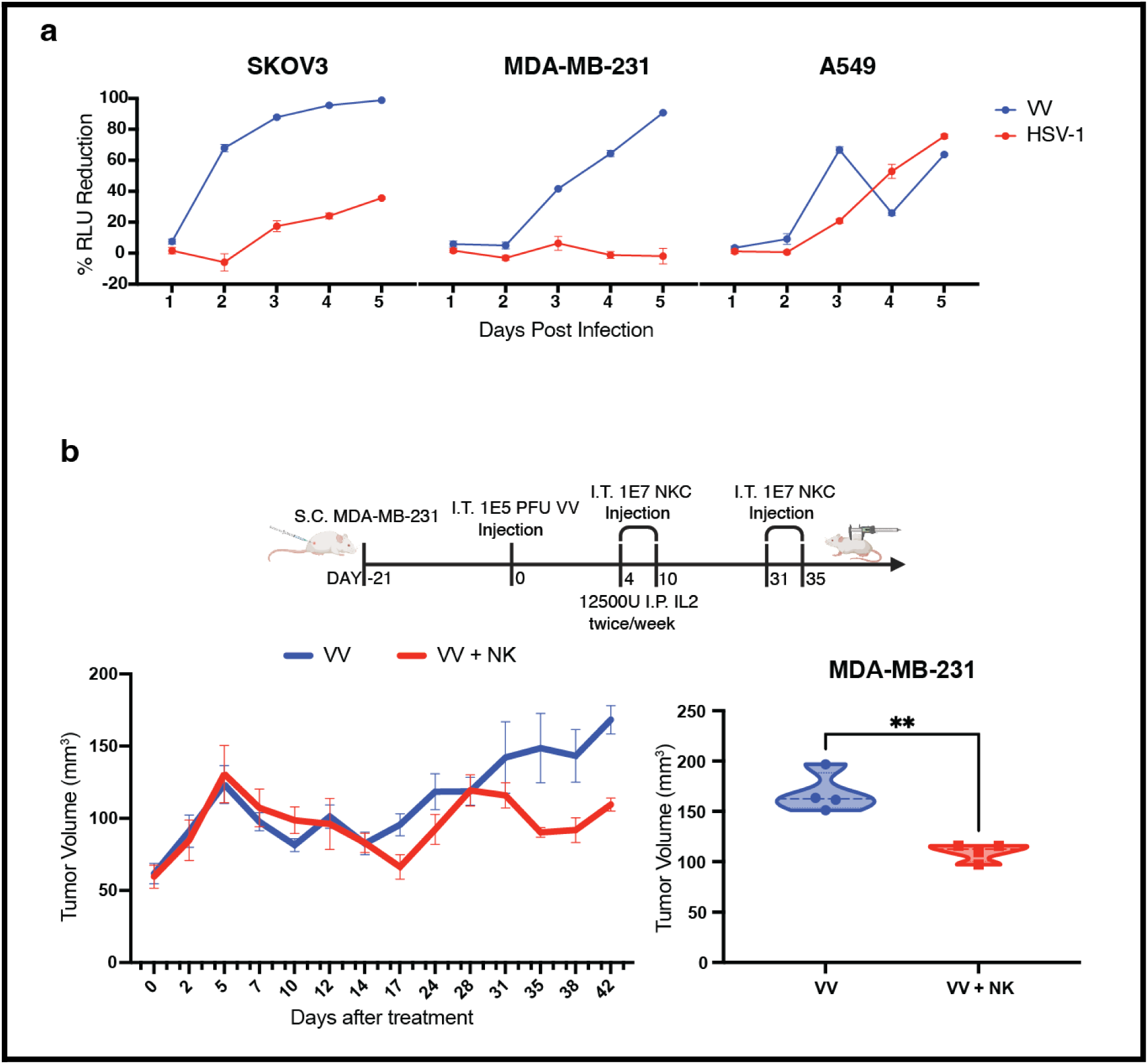
Longitudinal in vitro viral kinetics and long-term in vivo evaluation of vaccinia virus and NK cell combination therapy. (A) In vitro comparison of VV and HSV-1 cytotoxicity across SKOV3, MDA-MB-231, and A549 cell lines. CBR-expressing target cells were infected at an MOI of 0.01, and bioluminescence was measured daily for 5 days post-infection. Data represent the percentage reduction in relative light units (RLU) compared to untreated control wells at each 24 h interval (n = 4 replicates).(B) Extended in vivo evaluation of VV and EFT NK cell combination therapy in an MDA-MB-231 subcutaneous xenograft model. Top: Schematic of the animal study design showing tumor seeding (day -21), intratumoral (I.T.) injection of 1 x 10^5 PFU VV (day 0), and two cycles of I.T. EFT NK cell injections (1 x 10^7 cells per dose) on days 4/10 and 31/35. Mice were supplemented with IL-2 (12500 U, I.P.) twice weekly. Bottom Left: Longitudinal tumor volume measurements tracked through day 42 post-virus injection. Bottom Right: Violin plot comparing tumor volumes for the VV and VV plus NK cell groups at the day 42 endpoint. Data represent mean +/- SEM (n = 4 mice per group); P values were determined by an unpaired t-test (**).

**Fig. S5.**
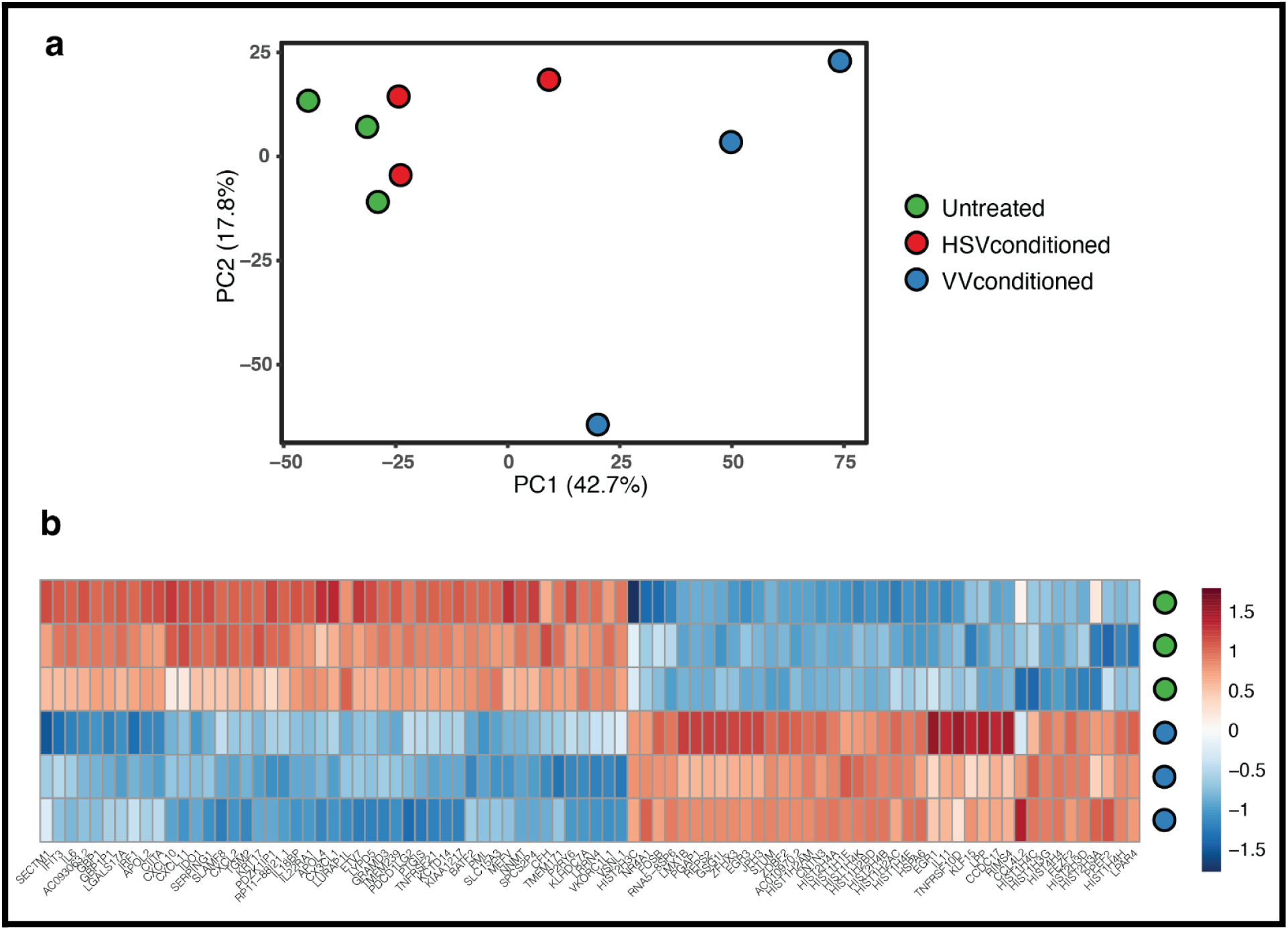
Principal component analysis and differential gene expression heatmap of conditioned media-treated EFT NK cells. (A) Principal Component Analysis (PCA) of EFT NK cells following treatment with untreated control media (green), HSV-1 conditioned media (red), or VV conditioned media (blue). The first two principal components, PC1 and PC2, account for 42.7% and 17.8% of the total variance, respectively. (B) Heatmap of differentially expressed genes in EFT NK cells across three independent biological donors. The top three rows represent samples treated with control media (C), and the bottom three rows represent samples treated with VV conditioned media (V). Sample identifiers correspond to specific biological donors (602, 607, 657, and 669). Color scale indicates the Z-score of normalized gene expression, ranging from -1.5 (blue) to 1.5 (red).

**Fig. S6.**
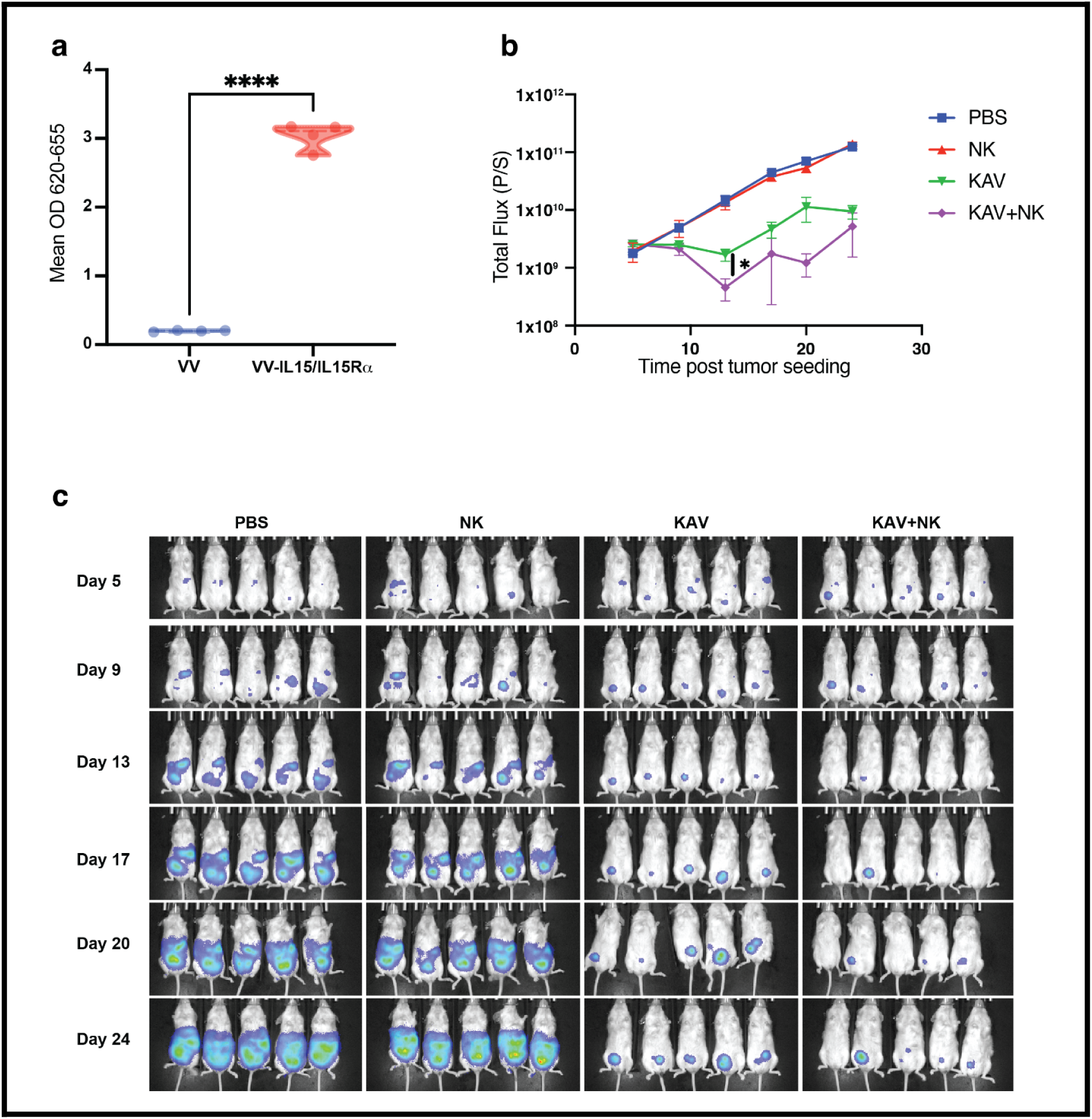
Functional validation of sIL15Ra-IL15 and bioluminescence monitoring of the SKOV3 intraperitoneal model. (A) Reporter assay for the functional validation of the sIL15Ra-IL15 fusion protein. Supernatants from A549 cells were collected 24 h post-infection with VV or VV-sIL15Ra-IL15 (MOI 0.5) and added to HEK293-Blue IL-2/IL-15 reporter cells. Functional activity is represented as Mean OD at 620-655 nm. Data represent mean +/- SEM (n = 4 replicates). (B) Longitudinal total flux (photons per second) measurements of luciferase-expressing SKOV3 tumors in an intraperitoneal xenograft model. Mice were seeded with SKOV3 cells on Day 0 and received an intraperitoneal (I.P.) injection of 1 x 10^5 PFU KAV or PBS on Day 6. EFT NK cells (1 x 10^7 cells per dose) were administered intravenously (I.V.) on Days 10 and 16 post-tumor seeding. (C) Representative bioluminescence images of mice from the indicated treatment groups on Day 5 (pre-treatment), 9, 13, 17, 20, and 24 post-tumor seeding.

**Fig. S7.**
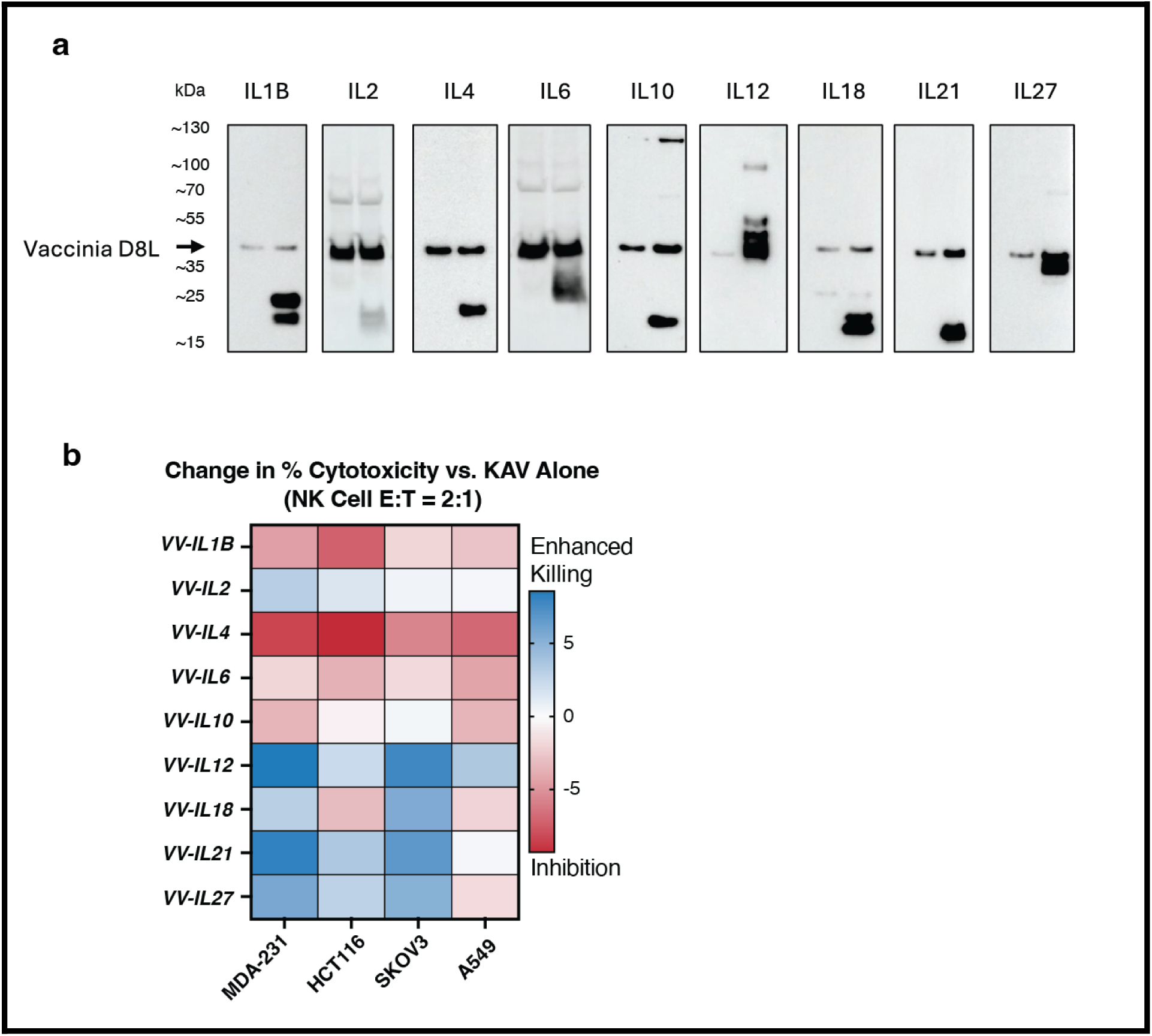
Biochemical validation and secondary combinatorial screening of the multi-cytokine KAV platform. (A) Western blot validation of cytokine expression. Side-by-side analysis of interleukin expression (IL-1B, IL-2, IL-4, IL-6, IL-10, IL-12, IL-18, IL-21, and IL-27) in A549 cell lysates 24 h post-infection with the respective vaccinia virus constructs. Viral D8L was used as a loading control for each blot. (B) Secondary 3D spheroid screening for candidate/supportive cytokine combinations.

**Fig. S8.**
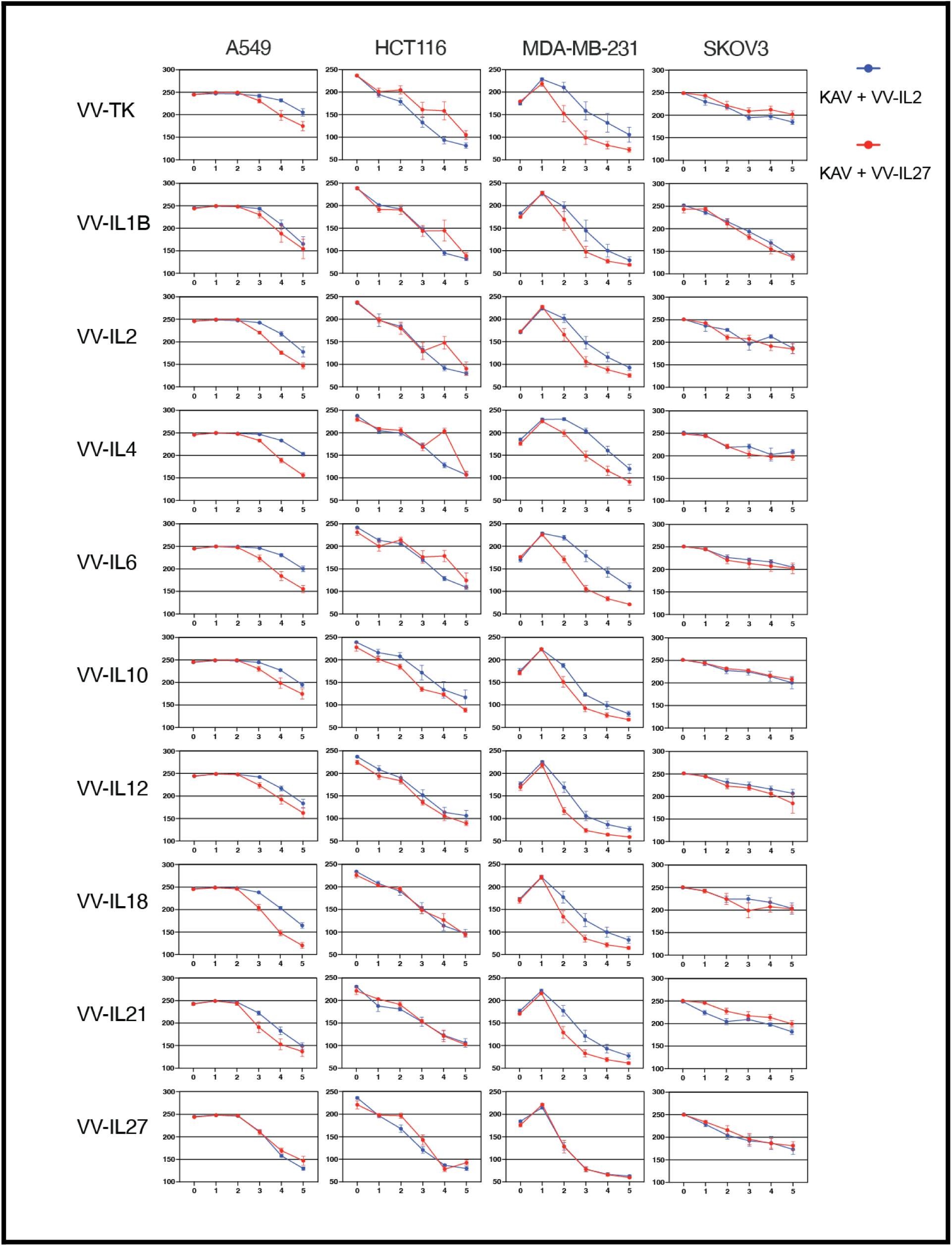
Combinatorial 3D spheroid screening of the multi-cytokine platform. Longitudinal GFP mean intensity of 3D tumor spheroids (A549, HCT116, MDA-MB-231, and SKOV3) treated with KAV plus VV-IL-2 (blue) or KAV plus VV-IL-27 (red) in combination with individual secondary interleukin-expressing viruses. Spheroids were infected at an MOI of 0.1 and cocultured with EFT NK cells at a 1:5 E:T ratio. Data represent mean +/- SEM from n = 4 independent biological donors.

**Fig. S9.**
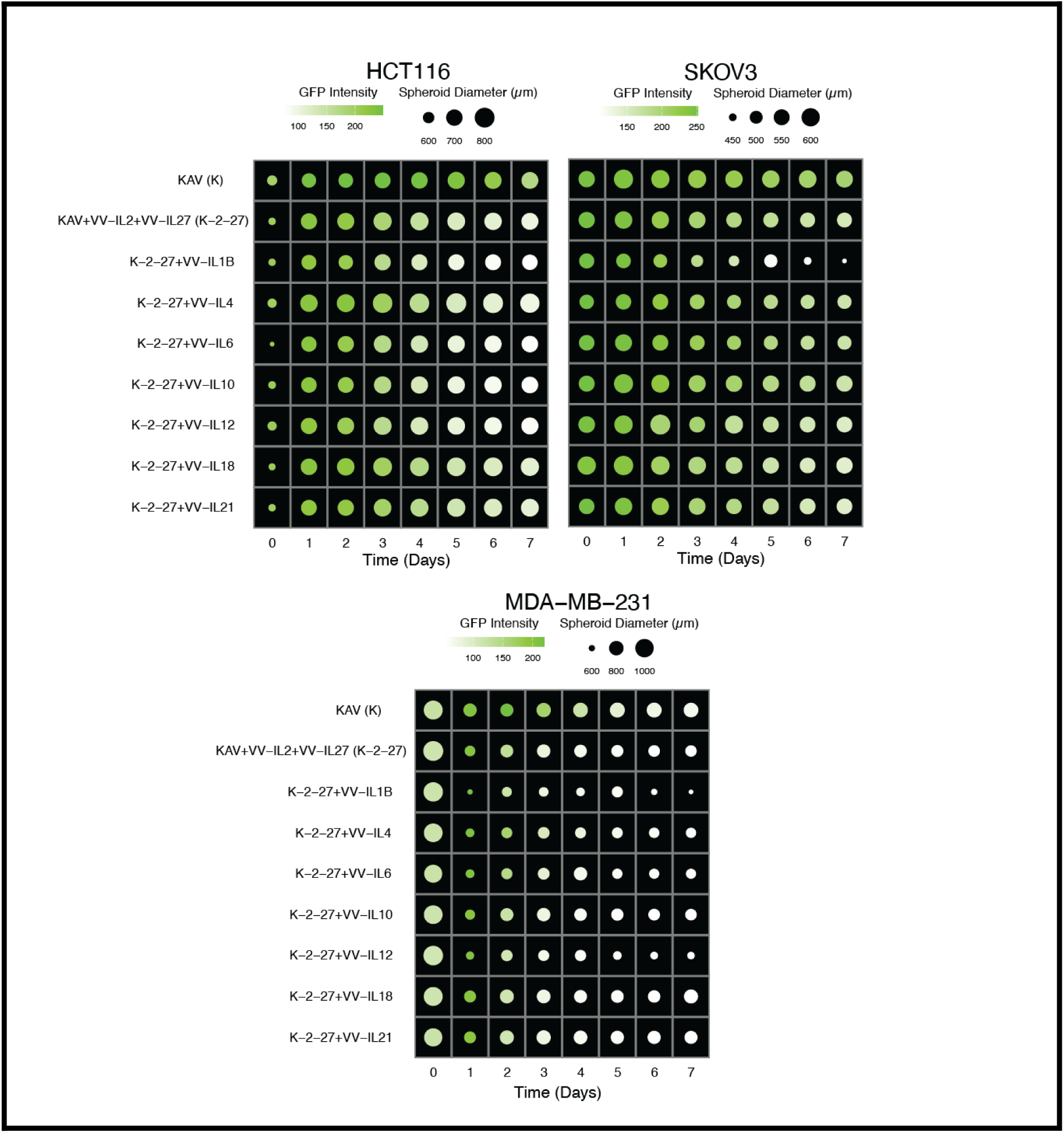
Simulation of 3D tumor spheroid growth and viability across multiple models. Dot plots representing average spheroid diameter (size) and mean GFP intensity (color) over 7 days post-treatment for HCT116, SKOV3, and MDA-MB-231 cell lines. Spheroids were treated with K (KAV), K-2-27 (KAV + VV-IL-2 + VV-IL-27), or the triple-cytokine combination with indicated interleukins IL-X (VV-IL-X) at a 1:10 E:T ratio.

**Fig. S10.**
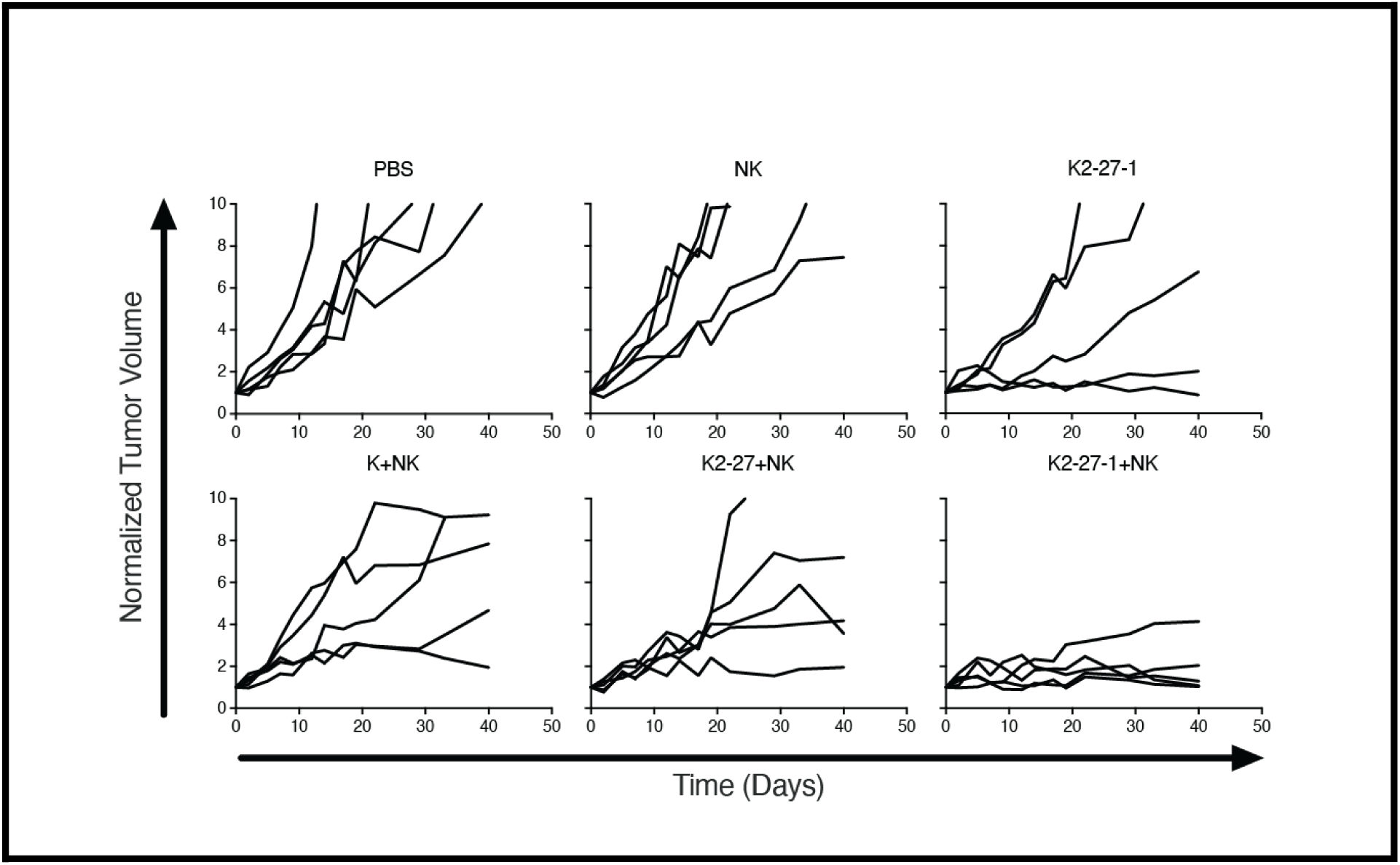
Individual tumor growth kinetics for the A549 subcutaneous xenograft model. Normalized longitudinal tumor volume for individual mice (n = 5 per group) from the study described in Figure 4E. Individual tracks represent the change in tumor volume relative to the initial size at the time of virus injection.

**Fig. S11.**
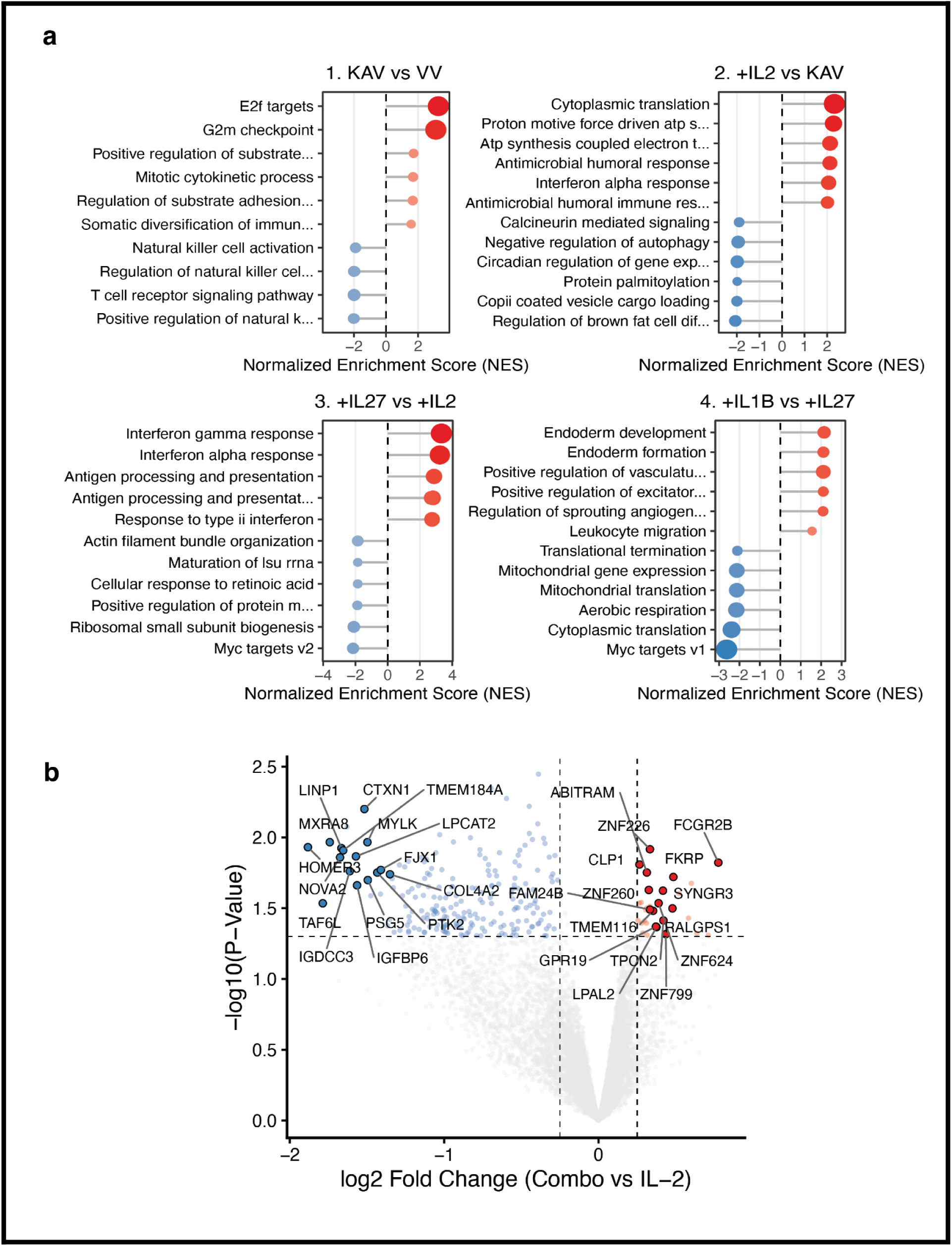
Transcriptomic signatures and GSEA of cytokine-primed EFT NK cells. Gene set enrichment analysis (GSEA) displaying normalized enrichment scores (NES) for sequential comparisons of viral supernatants: 1. KAV relative to VV, 2. (KAV +VV- IL-2) relative to KAV, 3. (KAV + VV-IL-2 + VV-IL-27) relative to (KAV + IL-2), and 4. (KAV + VV-IL-2 + VV-IL-27 + VV-IL-1β) relative to (KAV + VV-IL-2 + VV-IL-27). (B) Volcano plot showing differentially expressed genes between Combo (IL-2 + IL-1β) and IL-2 engineered NK cells; points represent individual genes with significant shifts in expression.

**Fig. S12.**
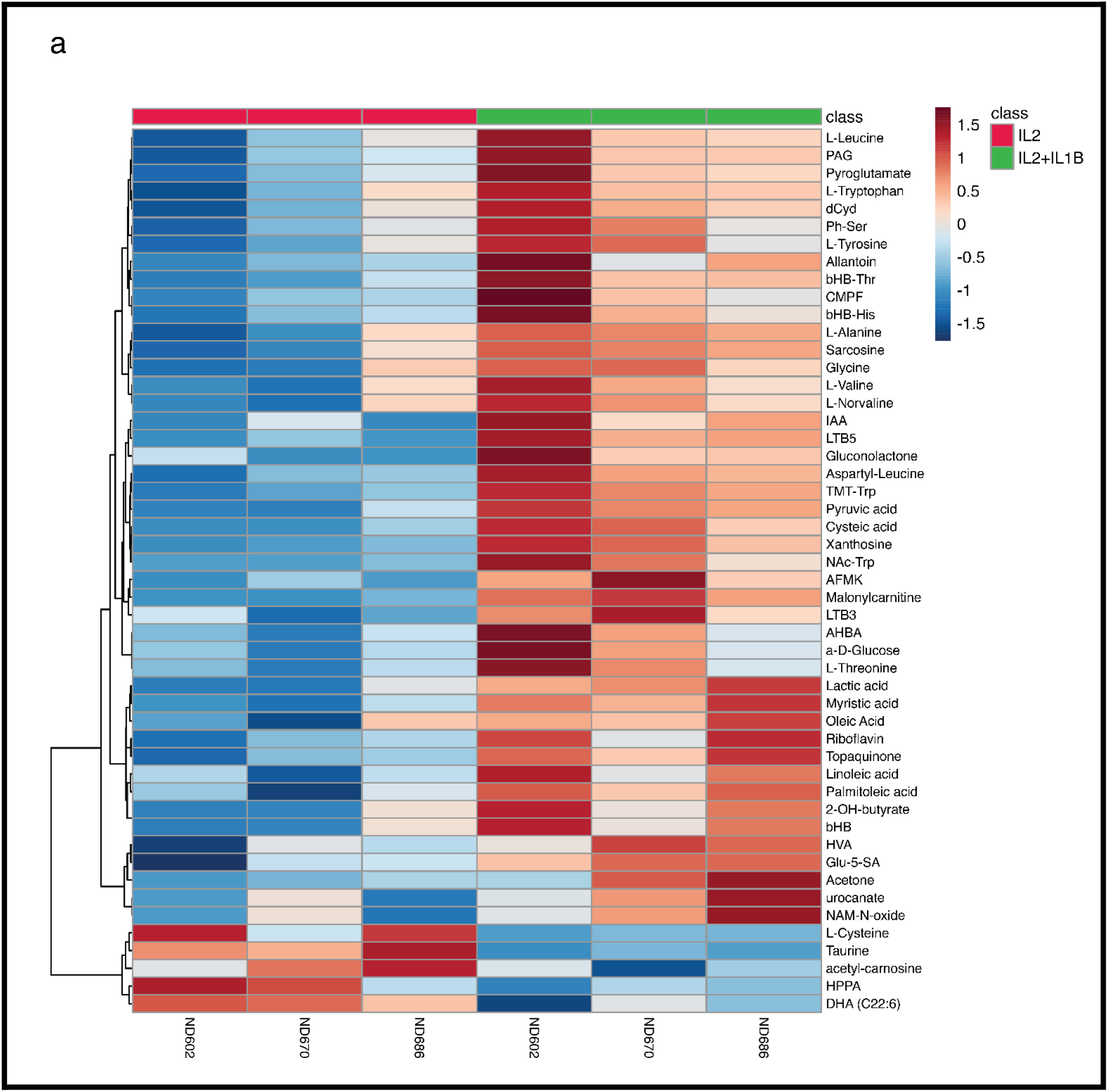
Comprehensive metabolomic profiling of cytokine-conditioned EFT NK cells. Heatmap of the top 50 differentially abundant metabolites between Combo (IL-2 + IL-1β) and IL-2 engineered NK cells identified via LC-HRMS. Metabolites were ranked by t-test p-value. Each column represents an individual healthy donor (n=3). Color indicates row-wise autoscaled (z-scored) relative abundance; rows were hierarchically clustered by Euclidean distance and Ward linkage.

**Fig. S13.**
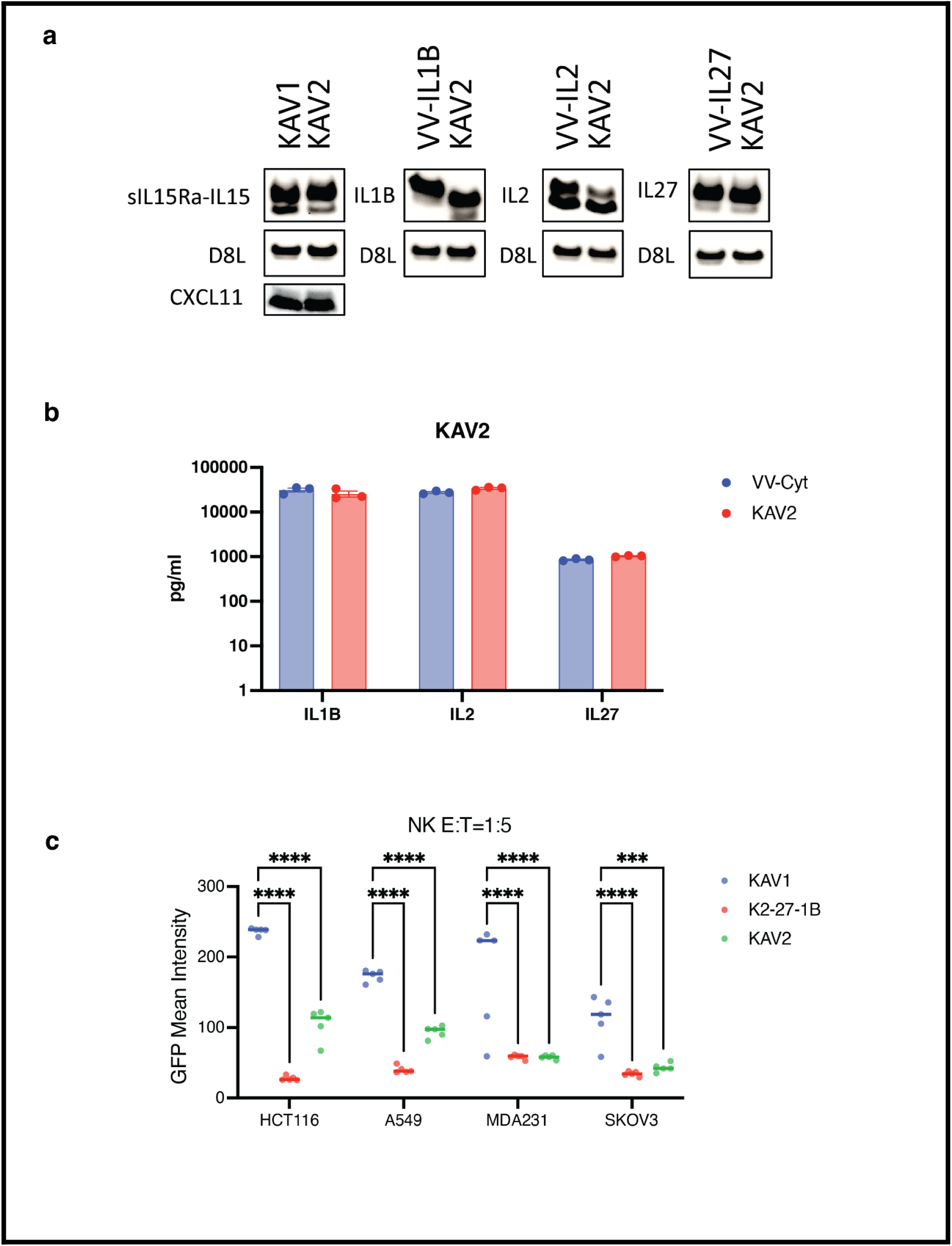
Validation of transgene expression and viral fitness for the consolidated KAV2 platform. (A) Western blot analysis of sIL-15Rα-IL-15, IL-1β, IL-2, IL-27, and CXCL11 in A549 cell lysates infected with KAV1, KAV2, or corresponding single-cytokine viruses. (B) ELISA quantification of secreted IL-1β, IL-2, and IL-27 in infected cell supernatants showing comparable expression levels between KAV2 and single-cytokine viruses. (C) 3D spheroid cytotoxicity of EFT NK cells (E:T 1:5) against indicated solid tumor lines following treatment with KAV1, the four-virus cocktail (K2-27-1B), or KAV2.

**Fig. S14.**
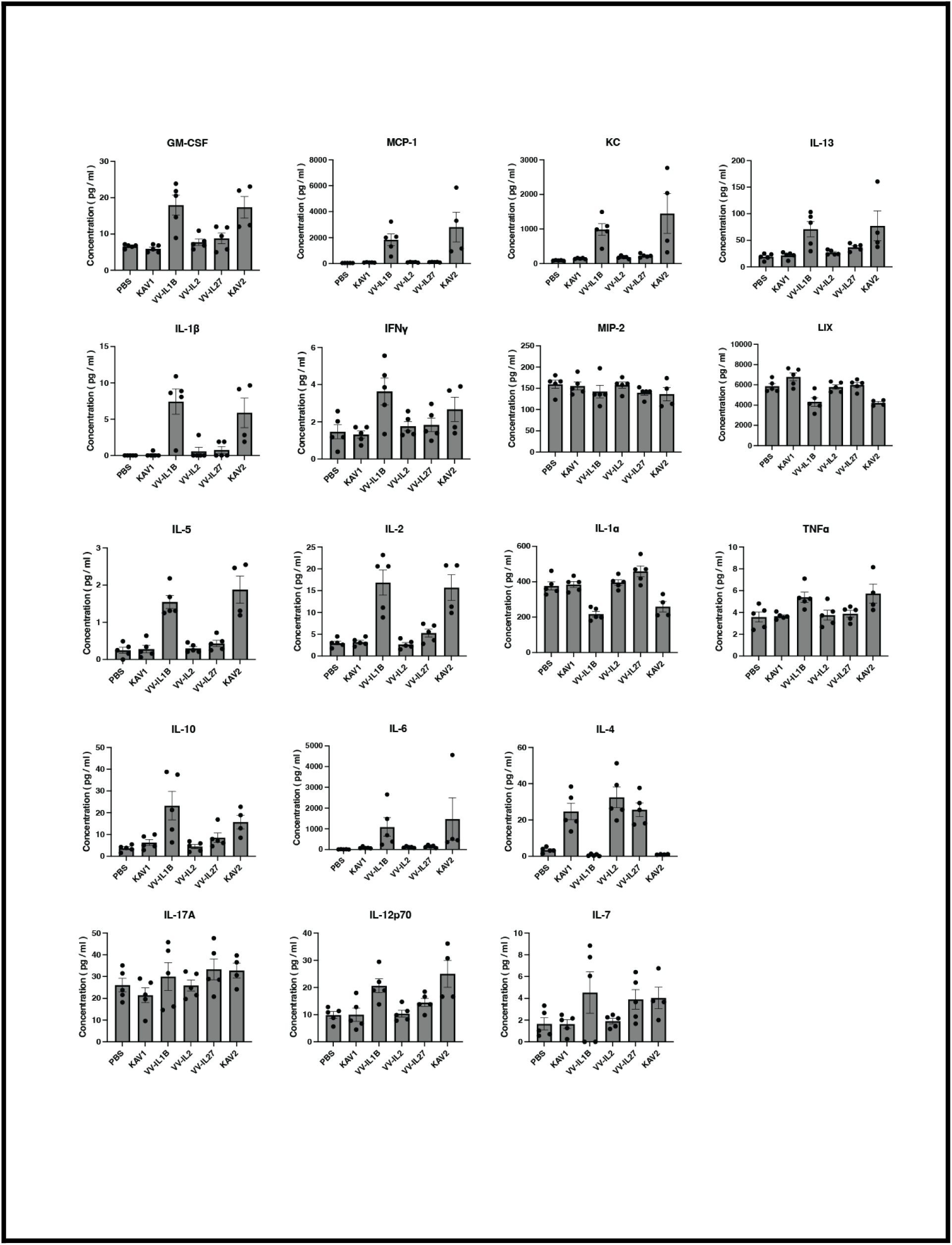
Quantitative characterization of the murine systemic cytokine response. Absolute concentrations (pg/mL) of 18 murine cytokines in the peripheral blood of NSG mice 3 days post-treatment with PBS, KAV1, VV-IL-1β, VV-IL-2, VV-IL-27, or KAV2. Data represent mean +/- SEM from n = 5 mice (except n=4 for KAV2) per group and were used to generate the Z-score radar plot in Figure 6I.

**Table. S1.** List of antibodies, fluorophore conjugates, and viability dyes used for flow cytometric analyses.

| Target Antigen | Fluorophore / Conjugate | Clone | Host / Isotype | Vendor / Supplier | Catalog Number |
|---|---|---|---|---|---|
| CD3 | BUV496 | UCHT1 | Mouse IgG1, κ | BD Biosciences | 612940 |
| CD3 | APC | HIT3a | Mouse IgG2a, κ | BD Biosciences | 555342 |
| CD56 (NCAM) | BUV737 | NCAM16.2 | Mouse IgG2b, κ | BD Biosciences | 612766 |
| CD8a | BUV496 | RPA-T8 | Mouse IgG1, κ | BD Biosciences | 612942 |
| CD159c (NKG2C) | BD RealYellow 586 (RY586) | 134591 | Mouse IgG1, κ | BD Biosciences | 753590 |
| CD159a (NKG2A) | PE-Vio 770 | REA110 | Recombinant human IgG1 | Miltenyi Biotec | 130-113-567 |
| CD158 (KIRs) | FITC | HP-MA4 | Mouse IgG2b, κ | BioLegend | 339504 |
| TIM-3 (CD366) | PE/Cyanine5 | F38-2E2 | Mouse IgG1, κ | BioLegend | 345051 |
| CXCR1 (CD181) | FITC | 8F1/CXCR1 | Mouse IgG2b, κ | BioLegend | 320606 |
| CXCR2 (CD182) | PerCP/Cy5.5 | 5E8/CXCR2 | Mouse IgG1, κ | BioLegend | 320718 |
| CXCR3 (CD183) | APC | G025H7 | Mouse IgG1, κ | BioLegend | 353708 |
| CXCR4 (CD184) | PE | 12G5 | Mouse IgG2a, κ | BioLegend | 306506 |
| Viability | Zombie NIR Fixable | N/A | Amine-reactive dye | BioLegend | 423105 |

